# Enhancing lipid production in plant cells through high-throughput genome editing and phenotyping via a scalable automated pipeline

**DOI:** 10.1101/2024.05.29.596527

**Authors:** Jia Dong, Seth W. Croslow, Stephan T. Lane, Daniel C. Castro, Jantana Blanford, Shuaizhen Zhou, Kiyoul Park, Steven Burgess, Mike Root, Edgar Cahoon, John Shanklin, Jonathan V. Sweedler, Huimin Zhao, Matthew E. Hudson

## Abstract

Plant bioengineering is a time-consuming and labor-intensive process, with no guarantee of achieving the desired trait. Here we report a fast, automated, scalable, high-throughput pipeline for plant bioengineering (FAST-PB). FAST-PB achieves gene cloning, genome editing, and product characterization by integrating automated biofoundry engineering of callus and protoplast cells with single cell matrix-assisted laser desorption/ionization mass spectrometry (MALDI-MS). We first demonstrate that FAST-PB can streamline the Golden Gate cloning process, with the capacity to construct 96 vectors in parallel. To prove the concept, using FAST-PB, we first found that PEG2050 significantly increases transfection efficiency by over 45%. To validate the pipeline, we established a reporter-gene-free method for CRISPR editing via mutation of *HCF136*, affecting cellular chlorophyll fluorescence. Next, we applied this pipeline for lipid production and found that diverse lipids were significantly enhanced up to sixfold through introducing multi-gene cassettes via CRISPR activation, and regenerated plant using this platform. Lastly, we harnessed FAST-PB to achieve high-throughput single-cell lipid profiling through the integration of MALDI-MS with the biofoundry, and differentiated engineered and unengineered cells using the single-cell lipidomics. These innovations massively increase the throughput of synthetic biology, genome editing, and metabolic engineering, and change what is possibly using single-cell metabolomics in plants.

## Introduction

Plant genetic engineering is needed more than ever to maintain global food security while battling ever-changing environmental conditions. However, currently engineering plants with desired traits is a complex and intricate process demanding significant time and labor (Mumm 2013). This process involves several key steps, including gene construct design, plant transformation or transfection, genome editing, and analysis and screening to identify the desired traits (Yin et al. 2017; Karlson et al. 2021). As a result, the development of engineered plants is typically a low-throughput and labor-intensive process, and represents one of the major limitations in plant biotechnology (Huang et al. 2022).

To overcome this limitation, we sought to integrate an automated biofoundry with single-cell metabolomics to expedite the engineering of plant genomes and characterization of cellular effects, which has never been attempted before. Biofoundries are specialized workstations that integrate robotics, high-throughput instrumentation, computer-aided design, and informatics to speed up iterative biological Design-Build-Test-Learn (DBTL) cycles in a scalable manner (Chao et al. 2017; Hillson et al. 2019; Zhang et al. 2021).

These workstations are highly reproducible, and optimize resource utilization by reducing human time and labor costs, increasing experimental throughput, and enabling researchers to devote more time towards experimental design as well as analysis and interpretation of results (Hillson et al. 2019). The Illinois Biological Foundry for Advanced Biomanufacturing (iBioFAB) has demonstrated success in automating synthetic biology processes such as plasmid assembly (Enghiad et al. 2022), yeast genome editing (Si et al. 2017), and antimicrobial discovery (Si et al. 2015). This success paves the way for the development of high-throughput plant genome editing technologies. Despite these possibilities, progress in automating plant biotechnology has been limited (Rigoulot et al. 2023).

Beyond developing engineered plants, a rapid and scalable method for characterizing genome editing is essential for development of a high throughput platform for plant improvement. Single-cell metabolomics is compatible with high throughput techniques, allowing the metabolic profiles of individual cells to be determined. Single-cell approaches allow the previously unknown phenotypic heterogeneity among cells to be investigated (Pandian et al. 2023). They also provide an immediate and dynamic snapshot of an individual cell’s functionality (Seydel 2021). Historically, metabolic data has been collected from cell populations, where average values lead to misleading interpretations about the state of each cell (Ali et al. 2019). Recent studies have shown that measurements of the average metabolome of a cell population conceals important information about the heterogeneity of individual cells, since cells are highly dynamic and constantly interact with each other and their environment (Lawson et al. 2015; Guo et al. 2021). Even two genetically identical cells often display different chemical metabolomes (Seydel 2021). Therefore, various bioanalytical tools have been developed for measuring single cell metabolomics, one example being matrix-assisted laser desorption/ionization mass spectrometry (MALDI–MS) (Neumann et al. 2019; Seydel 2021; Bourceau et al. 2023). These methods have recently been applied to lipidome profiling of single cells in the mouse brain (Zhang et al. 2023).

In this work, we have established a fast, <u>a</u>utomated, <u>s</u>calable, high-throughput pipeline for <u>p</u>lant <u>b</u>ioengineering (FAST-PB) (Figure 1). This pipeline seamlessly integrates the iBioFAB (Supplementary Figure S1) for automated synthetic biology with matrix-assisted laser desorption/ionization Fourier transform ion cyclotron resonance mass spectrometry (MALDI-FTICR-MS) for high-throughput single-cell lipid identification.

**Figure 1.**
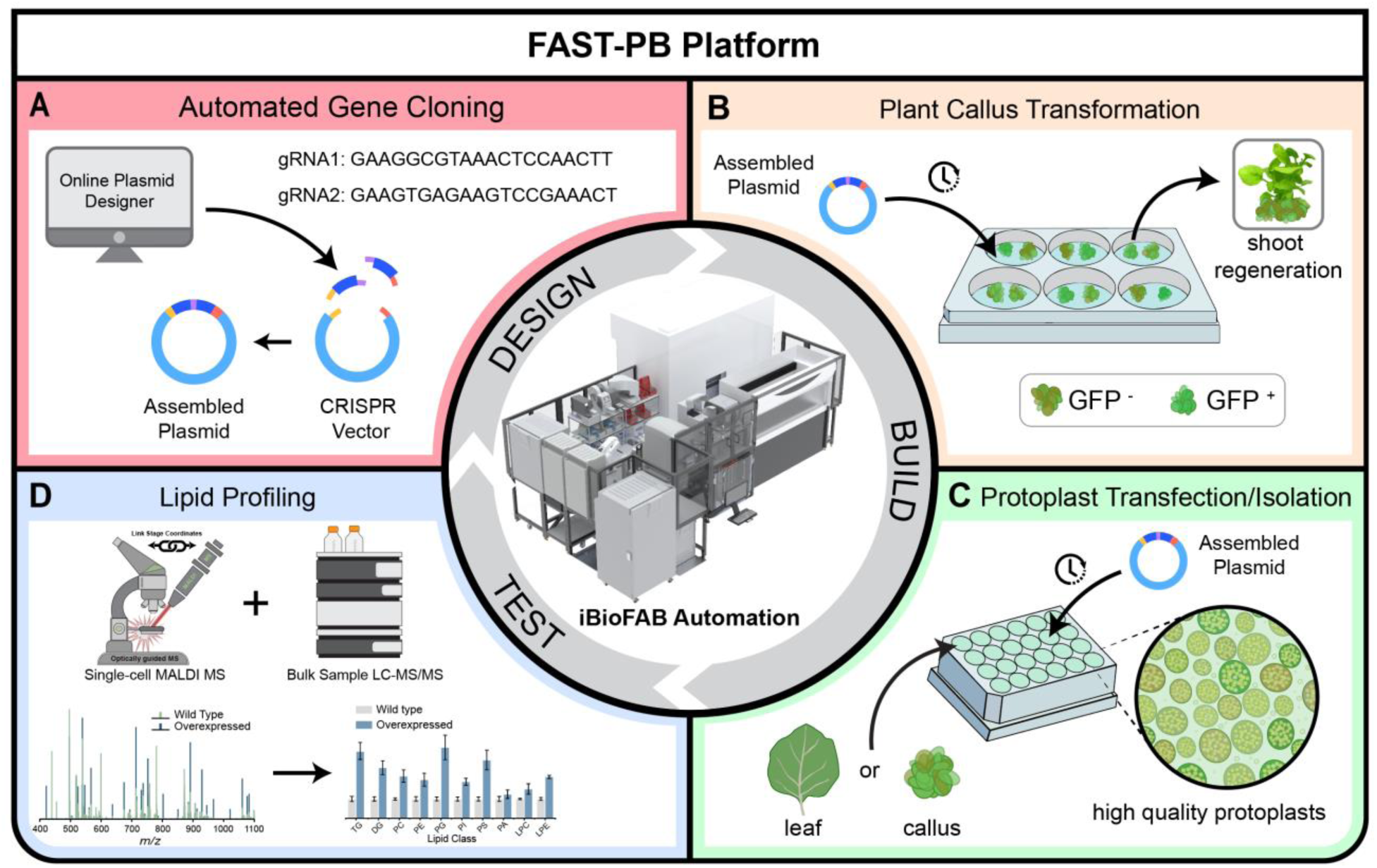
Overview of the FAST-PB for high-throughput genome editing and lipid engineering in protoplast and callus cell systems. (**A**) Automated gene cloning using the Golden Gate cloning method streamlines the process of plasmid assembly (Design). (**B**) Automated callus transformation with CRISPR vectors or strong promoter vectors to facilitate plant regeneration with the increased lipid production trait. GFP fluorescence indicates successful transformation (Build). (**C**) Automated protoplast isolation and transfection (Build). (**D**) Top: Protoplast cells from transgenic callus tissues applied on MALDI slides for lipid measurement and LC-MS analysis of lipids from callus and protoplast cells. Bottom: Characterization of lipids through MALDI spectra analysis and lipid class quantification (Test).

FAST-PB is compatible with not only protoplast cells that can be quickly isolated, transfected, and analyzed, but also callus cell cultures that can be maintained indefinitely for long-term studies and regenerated into a mature plant. As proof of concept, we applied this pipeline to assemble DNA constructs, engineer two cell factories (protoplast and callus) with a multi-gene stack giving a 2-6-fold improvement in lipid accumulation, and perform lipidomic analysis at the single-cell level in a high-throughput manner. Using this pipeline, we have several significant scientific findings: we first show that FAST-PB can be used for optimization of transfection efficiency, which can reach up to over 45%. Next, to quickly detect genome editing during the validation of our workflow, we developed a new reporter gene-free system. This system determines genome editing efficiency via the knockout of a photosynthetic gene, *HCF136*. This knockout leads to changes in chlorophyll fluorescence intensity in protoplast cells and the generation of slowly growing yellowish calli. Using this platform, we initially found that the lipids in protoplasts are stable over time, making them a useful platform for lipid engineering. Then we validated the function of two previously uncharacterized genes for lipid production, *DGAT1* and *WRI1* from *N. benthamiana* and maize. These two genes significantly increase the production of a variety of lipids and fatty acids by 2-6 fold through CRISPR activation system. Finally, our high-throughput single-cell lipid profiling finds the differentiation between transformed and untransformed cells, providing valuable information about how genetic engineering impact lipid metabolism at the cellular level. Overall, our pipeline has not only greatly accelerated transformation, CRISPR genome editing, single cell metabolomics, and plant regeneration, but also generated new scientific findings.

## Results

### Design of the FAST-PB pipeline

The iBioFAB is an integrated robotic platform that facilitates the development of fully automated workflows for scalable and high throughput synthetic biology applications. User-defined workflows are encoded into *Momentum* software, which coordinates communication between instruments, control of the robotic arm, and monitors location of the inventory of plates, samples, and consumables. FAST-PB is a *Momentum-*based biotechnology pipeline containing three main modules: (1) Design: CRISPR-P and Benchling online tools are used to design gRNAs and build digital CRISPR vectors containing Cas9 and gRNA sequences. A Golden Gate cloning workflow encoded into the iBioFAB utilizes a Beckman Coulter Echo acoustic liquid handler for preparation of PCR and DNA assembly reactions, an integrated TRobot2 thermocycler for reaction incubations, and a Tecan Fluent pipetting liquid handler for *Escherichia coli* transformations and plasmid extraction (Figure 1A, Supplementary Figures S1 and S2A, Supplementary Table S1). (2) Build: plasmids are transformed into *N. benthamiana* callus suspension cells (Figure 1B, Supplementary Figure S2B) or transfected into protoplast cells derived from maize or *N. benthamiana* leaves (Figure 1C, Supplementary Figures S2C-D). (3) Test: to complete the end-to-end automated trait optimization pipeline, two mass spectrometry methods are integrated for lipid profiling.

For single-cell lipid measurements using MALDI-FTICR-MS, callus-derived protoplasts are deposited onto indium tin oxide (ITO) coated microscopy slides. The locations of the single cells on the ITO-slide are then determined and probed using the MALDI laser for high-throughput lipid quantification. In addition, genome engineered plant cells can be analyzed via liquid chromatography tandem mass spectrometry (LC-MS/MS) (Figure 1D).

Overall, distinct plant cell systems exhibit unique advantages and disadvantages, with the selection of the optimal cell system often depending upon the specific application at hand. Protoplasts can be rapidly generated and easily chemically transfected but are generally short-lived, rendering them unsuitable for applications requiring complete plant tissues. On the other hand, callus systems are comprised of a mass of undifferentiated cells which can be maintained for longer periods and regenerated into plant organs. To develop a versatile platform applicable for the widest variety of plant engineering applications, we designed the FAST-PB platform (Figure 1) for compatibility with both callus tissue (Figure 1A, B, D) and protoplast cells (Figure 1A, C, D) as well as developing a custom technique that synergistically harnesses the benefits of both (Figure 1A-D).

### Automated protoplast isolation and transfection for scalable and rapid genome editing

Next, we established an automated workflow for high-throughput protoplast isolation, transfection, and gene mutant screening. Genetic engineering of plant cells is challenging because plant cell walls restrict the delivery of exogenous biomolecules (Demirer et al. 2019), and transformation and phenotypic screening is labor intensive (Lenaghan and Neal Stewart 2019; Squire et al. 2023). Protoplasts offer one potential solution due to their ease of production and transfection. Therefore, we removed plant cell walls to create protoplasts for plant genetic engineering.

The first step involved automating protoplast isolation from leaves of both maize and *N. benthamiana*. We translated the manual protocol for protoplast isolation (Supplementary Method 2) into an automated procedure (Supplementary Figure S2C). Briefly, leaf sections were excised and transferred into 24-well v-bottom plates for processing.

Samples were then digested, filtered, washed, and resuspended on the Tecan Fluent with required centrifugation steps performed by the F5 robotic arm and integrated centrifuge. Using this workflow, high quality protoplast cells were isolated from both etiolated maize and *N. benthamiana* leaves at a concentration of about 10^6^ cells / mL (Figure 2A-B).

**Figure 2.**
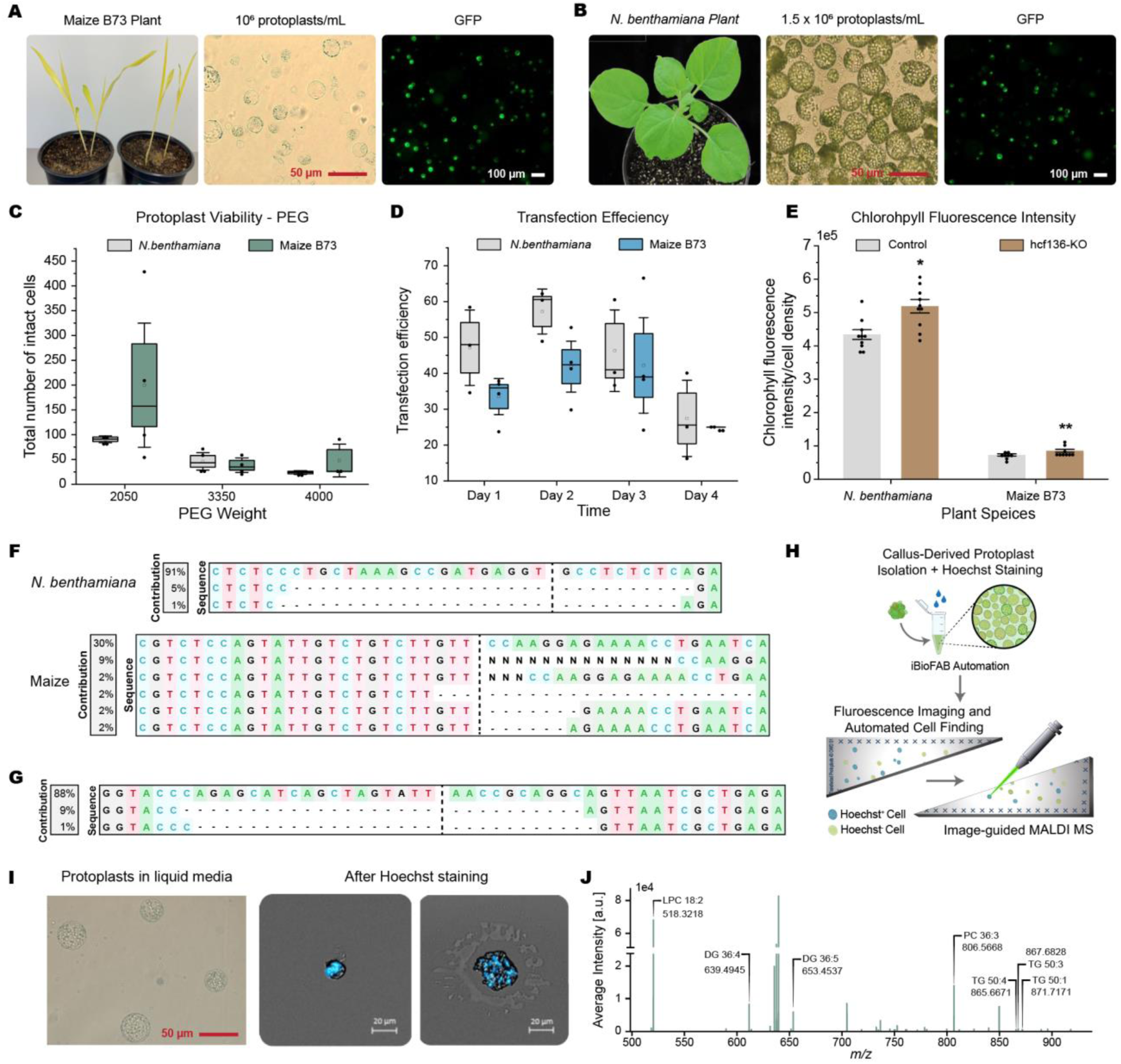
Automated protoplast isolation and transfection, automated callus transformation, detection of genome edits using *HCF136*, and single-cell lipid detection using MALDI MS. (**A**) Etiolated maize B73 (left) allowed successful protoplast isolation on the automation platform, yielding abundant high-quality protoplast cells (pictured at 40x magnification, middle). Subsequent transfection with the Cas9 vector (p201GFP-Cas9) induced GFP fluorescence in numerous cells (right). (**B**) Protoplast isolation from *N. benthamiana* leaves (left) using automation yielded abundant and high-quality protoplast cells (pictured at 40x magnification, middle). Protoplasts transfected with the Cas9 vector (p201GFP-Cas9) exhibit GFP fluorescence (right). (**C**) Total intact cell number was counted under three types of PEG (2050, 3350, 4000) treatments in the etiolated maize B73 and *N. benthamiana*. Biological replicate n = 4. (**D**) Transfection efficiency (Ratio of GFP-expressing and total intact cell numbers) in four days after transfection. Biological replicate n = 4. (**E**) Detection and quantification of chlorophyll fluorescence intensity following *HCF136* gene knockout in both maize and *N. benthamiana*. n = 10. (**F**) Mutation analysis of the *HCF136* gene performed using next-generation Sanger sequencing and Inference of CRISPR Editing (ICE) analysis (Hsiau et al. 2018; Dong et al. 2020) in maize and *N. benthamiana*. (**G**) Nucleotide sequences identified through ICE analysis following knockout of the *HCF136* gene in *N. benthamiana* callus. (**H**) Overview of the single-cell MALDI workflow for callus-derived protoplasts. (**I**) Protoplast cells in the W5 liquid media were imaged to check the quality before performing MALDI/MS (on the left). Then, using Hoechst staining method to stain the cells, they were imaged (In the middle and on the right). (**J**) Mass spectra with identified lipid species annotated. Error bars represent standard error. Asterisks indicate: \*\*\**P* < 0.001; \*\**P* < 0.01; \**P* < 0.05, calculated using two-tailed Welch’s t-test.

Next, we developed an automated workflow for plasmid transfection with protoplasts by encoding an established manual protocol (Supplementary Method 2) into *Momentum* (Supplementary Figure S2D). In general, plasmids, polyethylene glycol (PEG) transfection solution, MMG solution, and protoplast cells were mixed using the Tecan Fluent, followed by incubation. Subsequently, centrifugation was performed with the assistance of the F5 robotic arm and centrifuge. The washing and resuspension steps were then carried out using Tecan Fluent. We first applied this transfection workflow to optimize transfection efficiency by testing PEG with three different molecular weights: 2050, 3350, and 4000 g/mol. This choice is based on previous studies demonstrating that the molecular weight of PEG influences gene delivery efficiency (Zhang et al. 2008). Our experiments showed that PEG 2050 resulted in the highest fraction of GFP-positive cells (Supplementary Figures S3A-B) with the overall highest number of intact cells in both plants (Figure 2C and Supplementary Figures S3A-B), even though PEG 4000 is the most commonly used molecular weight for protoplast transfection in both *N.benthamiana* and maize. Next, we employed PEG 2050 to transfect cells to assess the viability of cells over time. We calculated the transfection stability over a period of four days by dividing the number of GFP-positive cells by the total number of intact cells. The largest proportion of cells expressing GFP was observed for the first two days, reaching up to 41% and 57% for maize and *N. benthamiana*, respectively (Figure 2D), but declined in the following days (Supplementary Figures S3C-D).

To demonstrate genome editing using automated protoplast isolation and transfection, we developed a reporter gene free cellular assay for loss of gene function. This endogenous genome editing assay is based on a single gene, *HCF136*, that increases chlorophyll fluorescence when its function is lost (Meurer et al. 1998). We used this assay to determine the effectiveness of the automated protoplast editing procedures using a simple plate reader assay for differential chlorophyll fluorescence. First, we isolated cells from both maize and *N. benthamiana* leaves then the two cell types were individually transfected with CRISPR knockout plasmids: A0502 (negative control) and A0502-HCF136 for *N. benthamiana,* and A1510 (negative control) and A1510-HCF136 for maize (Supplementary Table S1). Each plasmid transfection had 10 replicates, resulting in a total of 40 samples in a 96-well plate. Our results showed that we successfully induced mutations in *HCF136* in protoplast cells from both maize and *N. benthamiana* (Figure 2F), resulting in a significant enhancement in chlorophyll fluorescence intensity, as illustrated in Figure 2E. Targeted deletions in *HCF136* were confirmed via PCR, capillary sequencing and Inference of CRISPR Edits (ICE) analysis (Conant et al. 2022).

### Automated callus engineering to facilitate plant regeneration

In parallel, we developed an automated callus transformation workflow and integrated it into our FAST-PB pipeline (Supplementary Figure S2B). Unlike protoplasts, plant callus tissue is comprised of an unorganized mass of cells, which can be readily transformed and cultured for long periods (Ikeuchi et al. 2013). These properties make callus tissue a valuable tool for studying plant cell metabolomes, especially lipid metabolism (TSAI et al. 1982; Norouzi et al. 2022).

To initiate the callus transformation pipeline, three *Agrobacterium* GV3101 colonies harboring the A0502-HCF136 plasmid were selected using a colony picker integrated into the Tecan Fluent and inoculated into a 96-deep well plate containing selective media. The same procedure was performed for the control plasmid A0502. After overnight incubation, bacterial growth was quantified by measuring the absorption at 600nm (A600) using a Tecan Infinite plate reader. On the Tecan Fluent, 100 μL of overnight culture was transferred into a 6-well plate containing 3 mL of two-week-old callus suspension cells. After two-day co-culture, *Agrobacterium* was washed off using the Tecan Fluent. After one month of growth of calli on selective MS media, an automated genomic DNA extraction workflow (Supplementary Figure S2E) was employed to initiate mutant detection. We found that the *HCF136* gene contained targeted deletions after confirming mutants via capillary sequencing and ICE analysis. Additionally, we observed the mutated calli grew yellowish and unhealthy on the MS-selected media compared to the control (Figure 2G and Supplementary Figure S4A).

### Integration of single-cell MALDI MS into the FAST-PB pipeline for lipidomic analysis

To enable high-throughput lipid measurements and develop a complete end-to-end plant genome editing and cellular analysis pipeline, we harnessed the benefits of both protoplast and callus systems through a hybrid approach focused on single cell lipidomic analysis. We started by performing automated protoplast isolation from wildtype callus and then developed a single-cell MALDI-FTICR-MS lipidomic analysis pipeline for the isolated protoplasts (Figure 2H-I). Briefly, protoplasts are imaged in the W5 lipid media to check the quality of the cell before MALDI/MS (Figure 2I). Then cells are stained with a nuclear dye, deposited onto an indium tin oxide-coated microscopy slide, imaged under brightfield and fluorescence microscopy, then analyzed individually (Figure 2I). This technique enables automatic, quantitative and cost-effective single-cell lipid measurements with a high analytical throughput. As the first step, we conducted traditional LC-MS/MS lipid profile measurements on bulk isolated protoplasts derived from wild type calli, to ensure the consistency of the datasets obtained from the LC-MS/MS and MALDI-MS. Of the detected lipid types from MALDI-MS, 33% were matched to the LC-MS/MS lipid datasets. Using this workflow, MALDI spectral analysis revealed a variety of lipid types (Figure 2J). All these results indicate that MALDI-MS has the capability to detect a wide range of lipids in individual cells.

### Harnessing the FAST-PB pipeline to generate cultivars with enhanced lipid production

Lipids in plants serve multiple functions, including energy storage, structural support, protection, and signaling (Okazaki and Saito 2014; Xie et al. 2021). To effectively explore lipid metabolism using protoplasts, it is essential that lipids remain relatively stable throughout the experimental period. Thus, we applied our automated protoplast isolation workflow to measure the lipid content in protoplasts over a four-day period.

Apart from a substantial increase in triacylglyceride (TAG) content in *N. benthamiana* over three days, most lipids remained stable for four days in both plants (Figure 3A-B).

**Figure 3.**
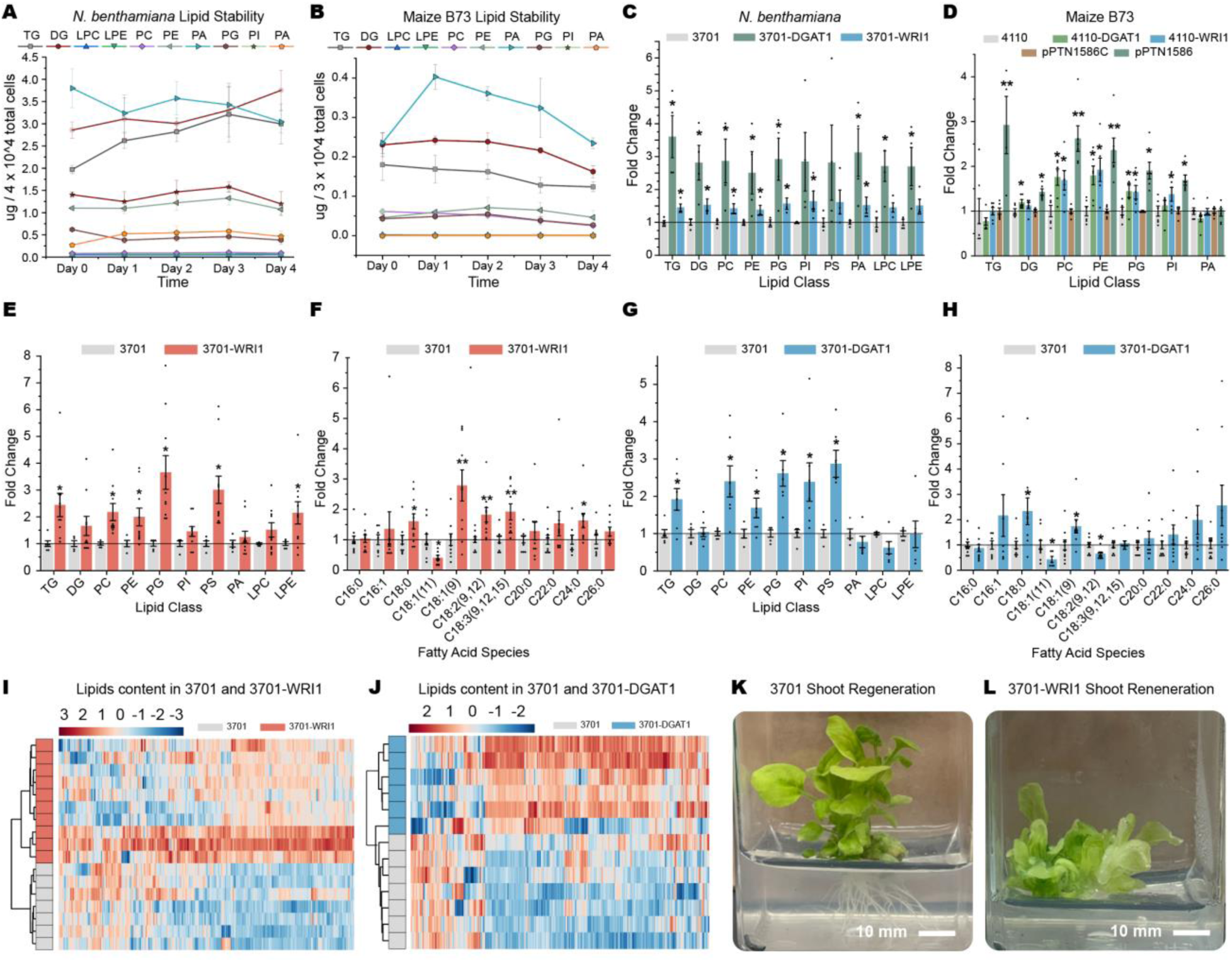
Lipid profiling and comparative analysis in protoplast cells and genetically engineered calli using LC-MS/MS data. (**A-B**) Lipid classes were measured across four days using LC-MS/MS in *N. benthamiana* (**A**) and etiolated maize B73 (**B**). Biological replicate n = 6. (**C**) Lipid analysis and quantification after overexpression of 3701-WRI1 and 3701-DGAT, with 3701 as a control in *N. benthamiana*. Biological replicate n = 4. (**D**) Lipid analysis and quantification after overexpression of 4110-WRI1 and 4110-DGAT, pPTN1586, with 4110 and pPTN1586C as controls in etiolated maize B73. Biological replicate n = 5. In each heatmap, the clustered grey bar at the top represents the control group, while the red or blue bar indicates the overexpression genes. n >= 6. (**E** and **G**) Quantification of lipid classes through fold change calculation for the overexpression of the *WRI1* gene (**E**), and overexpression of the *DGAT1* gene (**G**). n >= 5. (**F** and **H**) Fatty acids quantification through fold change formula for the overexpression of *WRI1* gene (**F**), the *DGAT1* gene (**H**). n >= 5. (**I-J**) Heatmaps comparing control and experimental samples for lipid data classification and statistical analysis for the overexpression of the *WRI1* gene (**I**), and an overexpression of the *DGAT1* gene (**J**). (**K-L**) Top (**K**): Shoot and root regeneration from *N. benthamiana* calli carrying an empty vector 3701. Bottom (**L**): Shoot regeneration from genetically engineered calli (3701-WRI1: Overexpression of *WRI1* through CRISPR activation). Error bars represent standard error. Asterisks indicate: \*\*\**P* < 0.001; \*\**P* < 0.01; \**P* < 0.05, calculated using two-tailed Welch’s t-test.

We then utilized the FAST-PB pipeline to investigate the effects of overexpressing *WRI1*, *DGAT1* and *Oleosin* on the lipidomic profile of maize and *N. benthamiana* protoplast cells using CRISPR activation or strong promoter systems. *N. benthamiana* protoplast cells were transfected with P-A3701, P-A3701-DGAT1, P-A3701-WRI1, and p201GFP-Cas9, while maize protoplast cells received separate transfections with A4110, A4110-DGAT1, A4110-WRI1, pPTN1586C, pPTN1586, A1510, A1510-HCF136, and p201GFP-Cas9.

In *N. benthamiana*, overexpression of *WRI1* via CRISPR activation increased accumulation of various lipid classes while overexpression of *DGAT1* significantly enhanced TAG content (Figure 3C). In maize, overexpression of either *WRI1* or *DGAT1* through CRISPR activation also resulted in increased accumulation of many types of lipids (Figure 3D). The same trend was found after overexpression of all three *Oleosin*, *WRI1*, and *DGAT1* genes (OWD) using a strong promoter (Figure 3D). In addition, we confirmed *DGAT1* was overexpressed through CRISPR activation in maize cells (Supplementary Figure S4C). These findings demonstrate a complete end-to-end automated pipeline for implementing genetic perturbations and characterizing lipid profiles using protoplast cells to accelerate the plant genetic engineering process.

Following this, we applied the FAST-PB pipeline to callus cells to assess two distinct lipid genes. We transformed C-A3701, C-A3701-DGAT1, and C-A3701-WRI1 into callus suspension cells with two replicates for each transformation event. We individually overexpressed *DGAT1* and *WRI1* using CRISPR activation (Supplementary Figure S4B) and confirmed that both genes were overexpressed by RT-qPCR (Supplementary Figure S4D). Overexpression of *WRI1* increased accumulation of a variety of lipids and fatty acids (Figure 3E-F) and overexpression of *DGAT1* induced a similar overall profile (Figure 3G-H), showing that CRISPR activation of lipid metabolic genes altered lipid composition. Individual overexpression of either *WRI1* (Figure 3I) or *DGAT1* (Figure 3J) decreased accumulation of some lipids, although a far larger number either increased or remained unchanged. After confirming an increase of lipid content in these genetically engineered calli, we initiated the process of plant regeneration. After growth in MS induction media, plants were regenerated from engineered calli (Figure 3K-L), demonstrating that the FAST-PB pipeline enables designing, building, and testing plant genetic engineering using robotics before regenerating whole plants.

### Application of the FAST-PB pipeline onto high-throughput single-cell lipid profiling to accelerate plant engineering

The FAST-PB pipeline was used to transform plasmids pMDC43 and pMDC43-OWD into callus suspension cells (Figure 4A-B) and isolate protoplasts from the transgenic calli. Callus cultures transformed with either an *Oleosin*, *WRI1*, and *DGAT1* (*OWD*) overexpression cassette or an empty vector control were confirmed by observation of GFP fluorescence and then stably maintained on selective agar plates (Figure 4A-B). The lipids and fatty acids in these genetically engineered calli were analyzed from tissue samples with LC-MS/MS. Compared to control cells, overexpression of *OWD* led to increased accumulation of many lipids and fatty acid species (Figure 4C), indicating overall increased cellular lipid accumulation.

**Figure 4.**
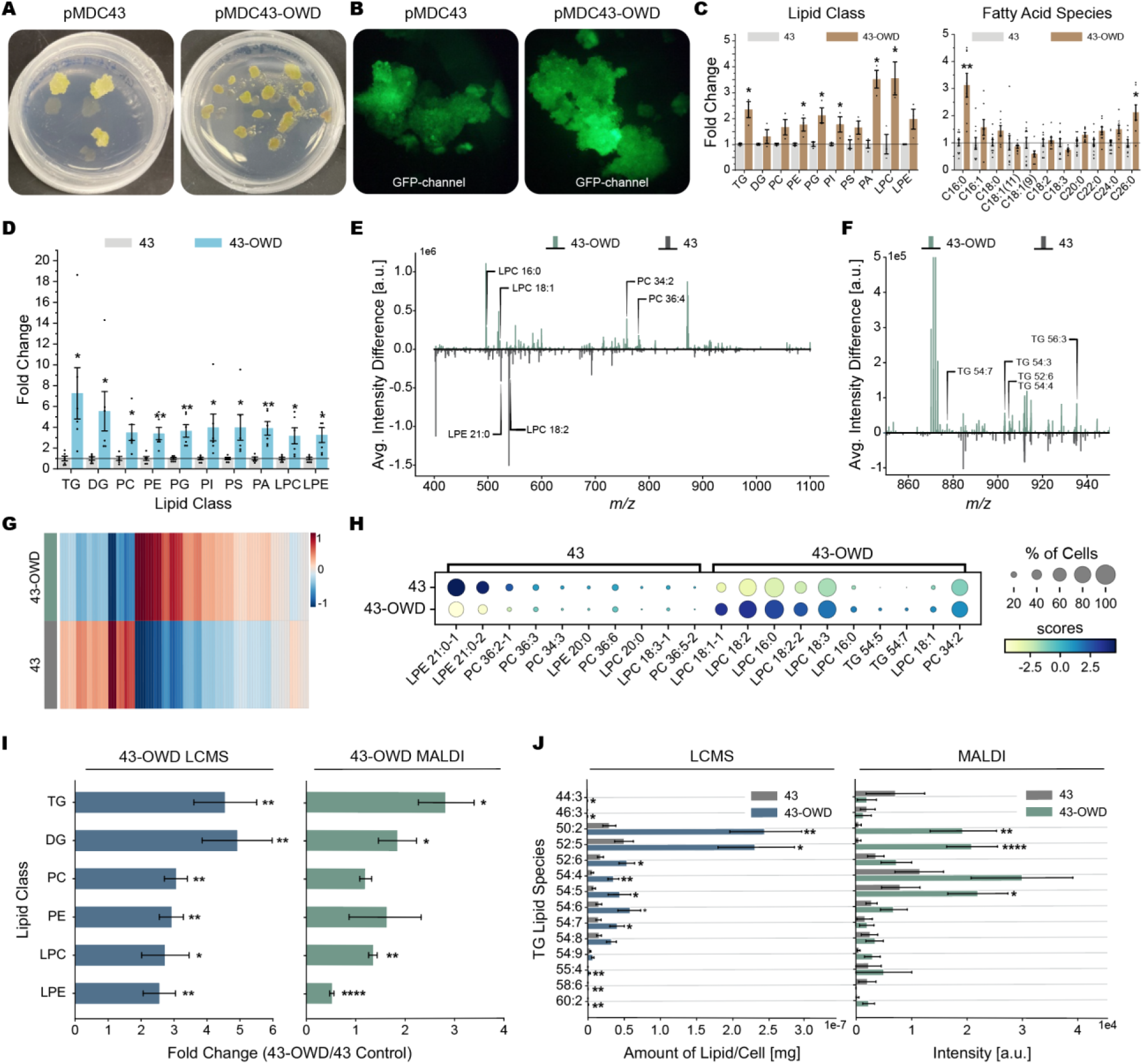
Lipidomic analysis of genetically engineered protoplasts samples using LC-MS/MS and high-throughput single-cell measurements using MALDI FT-ICR MS. (**A-B**) Bioengineered calli overexpressing *Oleosin, WRI1, and DGAT1* (OWD) genes with a strong 35S promoter grown on MS media with hygromycin resistance (**A**) and displaying strong GFP fluorescence in the calli tissue (**B**). (**C**) Left: Lipid profiling in genetically engineered calli. Right: The comparison of fatty acid species in bioengineered calli. (**D**) Lipid profiling in callus-derived protoplast cells. (**E**) Subtracted average mass spectra for control (43) and genetically overexpressed (43-OWD) protoplasts across the entire lipid range. The accurate *m/z* values for can be found in *SI Appendix,* Supplementary Table S3. (**F**) Selected *m/z* window around the TAG lipid region, with detected lipid species annotated. The accurate *m/z* values for can be found in Supplementary Table S3. (**G**) Heatmap of all the LCMS-matched lipid features from the single-cell MALDI dataset. (**H**) Ranked dot plot showing the top ten lipid features for the control (43) and genetically engineered (43-OWD) protoplast samples. Samples are colored by their p-value-modulated z score and the size of each dot represents the fraction of the total samples that had each feature. (**I**) Comparison of lipid class between the LC-MS/MS and MALDI datasets. (**J**) The comparison of TAG lipid features between LC-MS/MS and single-cell MALDI detection. Biological replicate n > = 3 for all MALDI and LC-MS/MS datasets. Error bars represent standard error. Asterisks indicate: \*\*\*\**P* < 0.0001; \*\*\**P* < 0.001; \*\**P* < 0.01; \**P* < 0.05, LCMS statistics were calculated using a two-tailed Welch’s t-test and MALDI data was analyzed using a two-tailed Mann-Whitney test.

To assess the consistency of lipid profiles between genetically engineered calli and protoplast cells derived from these calli, we directly compared these platforms using bulk LC-MS/MS analysis. Consistent with the results obtained from extracting lipids from engineered calli, the callus-derived protoplasts exhibited increased lipid accumulation after overexpression of either *WRI1* or *DGAT1* or simultaneous overexpression of the three-gene *OWD* stack (Figure 4D and Supplementary Figure S5), demonstrating that lipid profiles of callus-derived protoplasts represent the original callus.

High-throughput single-cell lipid metabolite profiling was then performed on the callus-derived protoplasts using MALDI-MS. Single-cell MALDI data was mass matched to the LC-MS/MS datasets for putative lipid assignment (Supplementary Table S3). MALDI spectral analysis revealed higher levels of several lipid classes after overexpression with 43-OWD when compared against the negative control (Figure 4E-F). A heatmap overview of the trends of all identified lipid species shows a similar increase across most lipid classes (Figure 4G). Utilizing the MALDI dataset, we observed a significant increase in the accumulation of various lipid classes, particularly storage lipid TAGs, in response to *OWD* expression (Figure 4H). These findings indicate that MALDI analysis of protoplasts is a promising method for single-cell lipid measurements in plants.

To ensure the accuracy and reliability of the MALDI method for lipid measurement, we compared single-cell MALDI to the reference method, LC-MS/MS on bulk cells. We found a high degree of consistency between the two datasets, both in terms of lipid classes and species-level trends, such as TAGs (Figure 4I-J). This finding indicates that single protoplast MALDI-FTICR-MS facilitates accurate and efficient high-throughput lipid measurement on single plant cells, positioning it as a valuable alternative to the LC-MS/MS approach. Similar results on the comparability and concordance between LC-MS and MALDI have also been recently reported in human cells (Martín-Saiz et al. 2023). Moreover, our single cell measurements and data processing have the capacity to segregate the engineered and unengineered cell types within the 43-OWD group, demonstrating that based on single-cell heterogeneity, we can estimate which protoplasts have been genetically engineered (Supplementary Figure S5E).

Taken together, by coupling callus transformation with protoplast isolation and MALDI-FTICR-MS analysis, FAST-PB embodies a high-throughput, automation-friendly, end-to-end pipeline for plant genetic engineering and characterization at the single cell level.

This can be used for investigating the effects of specific genetic perturbations on lipid metabolism, and potentially other metabolic pathways.

## Discussion

Traditional plant genetic engineering involves labor-intensive, low-throughput processes encompassing gene construction, transfection or transformation, genome editing, mutant identification, gene function analysis, and plant regeneration (Mumm 2013; Yin et al. 2017; Karlson et al. 2021). To automate these steps and achieve high throughput, we successfully established the FAST-PB pipeline comprising automated gene constructions, protoplast cell isolation and transfection, callus transformation, genomic DNA extraction, and lipid profiling. Our results demonstrate that the FAST-PB pipeline is highly applicable and scalable for engineering plant cell traits through the combination of automation, transformation, editing and metabolomics, and represents a key step towards fully automated plant biotechnology.

While biofoundries excel in precision, scalability, and cost-efficiency for genome editing and metabolic engineering (Si et al. 2017), they have yet to be applied to plant synthetic biology. Here we fill this gap with our broadly applicable and scalable plant genetic engineering pipeline. Our automated protoplast engineering workflow enables rapid isolation and transfection of up to 96 individual samples in a short amount of time.

Another potential benefit of this workflow is the ability to simultaneously optimize multiple parameters (e.g. PEG types, mannitol concentrations, and incubation times) for transfection efficiency. As an example, we were able to quickly optimize transfection efficiency using three PEG batches with different average molecular weights (2050, 3350, and 4000) on the iBioFAB platform, finding that the PEG 2050 consistently enhanced transfection efficiency while preserving the integrity of the protoplast cultures (Figure 2C). Furthermore, we envision applications of our automated protoplast engineering workflow for high-throughput assessments of in vivo gene editing efficiency of specific guide RNAs, subcellular localization experiments via fused fluorescent proteins, or confirmation of protein-protein interactions through bimolecular fluorescence complementation assays. Our automated callus transformation workflow enables simultaneous processing of up to twelve six-well plates at a time for a total throughput of 96 samples and facilitating plant regeneration (Figure 3K-L), opening the door to high-throughput screening of genes related to whole-plant traits such as plant pathology, biofuel production, and crop yield improvement.

Single-cell technology has gained significant attention in recent years (Roy et al. 2018). Plant tissues encompass a multitude of distinct cell types, each assuming specialized roles, exhibiting diverse molecular behaviors, and generating unique metabolites in response to biotic and abiotic stresses (Cole et al. 2021). Profiling whole plant tissues using traditional bulk methods can obscure and dilute signals associated with specific single cells. However, plant cell walls and tissue structures tend to complicate single cell analysis. Therefore, we automated the isolation of protoplast cells using the iBioFAB platform, followed by measurement of lipid profiles in each single cell using MALDI/MS technology to achieve a high-throughput assay, and validation of the results in comparison to low-throughput LC-MS/MS assays on bulk cells. We were able to measure a wide variety of lipid classes in single cells, including storage lipids (TAG and DAG) and phospholipids (PC, PE, LPC, and LPE) (Figure 4). Therefore, we successfully established an automated platform for high-throughput genome editing and lipid engineering with single-cell resolution.

Our platform and single-cell studies in general have a variety of potential applications. Previous studies have identified specific cell types related to processes such as oil production (Taylor et al. 2021), plant pathogen responses, and environmental stress (Cole et al. 2021). Our platform enables quickly transfecting cells for high-throughput gene analysis and enables dissection of gene function at the single cell level. Moreover, our workflows can accelerate genomic and metabolic engineering processes within cell factories, facilitating plant transformation and regeneration. Lastly, within each cell, various types of organelles / vesicles play a role in disease or stress tolerance (Cui et al. 2020; Urzì et al. 2021). Recent reports demonstrate the ability of MALDI MS to probe these subcellular structures (Castro et al. 2021, 2023; Eberwine et al. 2023), although not all these methods have yet been applied to characterizing plant vesicles. The ability to examine the contents of these subcellular structures, an obvious next step for our platform, could deepen our understanding of lipid function in plant health and diseases.

Lipid engineering in plants is gaining importance in agriculture as lipids play essential roles as energy storage, cell membrane components, and signaling molecules in plant growth and defense mechanisms (Raczyk and Rudzińska 2015; Mamode Cassim and Mongrand 2019). In this study, we focused on three lipid-related genes (*WRI1*, *DGAT1* and *Oloesin*) and employed a classic push-pull-protection strategy (Vanhercke et al. 2014, 2019; Volk et al. 2023) to enhance lipid production through lipid engineering in cell systems. Following the overexpression of these three genes either individually or in a triple combination, using either the traditional strong promoter system or the more versatile CRISPR activation method, we observed a substantial increase in the accumulation of various types of fatty acids and lipids (Figures. 3-4). The results indicate that our cell systems, including protoplasts and callus suspension cells, as well as our three automated workflows, are highly suitable for lipid metabolic engineering. In addition, besides these three genes, numerous other genes are involved in oil production in leaves (Vanhercke et al. 2014). Furthermore, lipids play a significant role in interactions with various pathogens and microbes, serving as mediators in the communication between hosts and pathogens (Raczyk and Rudzińska 2015). Hence, we have the potential to stack or combine additional genes from these three steps (pull, push, and protection) with the aim of enhancing vegetative lipid production for bioenergy, or to explore the role of lipid species in plant disease resistance pathways to facilitate high-throughput engineering of disease resistance or microbial symbiosis traits in crops.

In summary, the FAST-PB pipeline developed here has potential to transform genome editing, metabolic engineering, and metabolite profiling in plants by expanding the toolkit for trait discovery and manipulation by executing iterative design-build-test-learn cycles in small cell cultures. This workflow streamlines discovery, characterization, and fine-tuning of traits in highly scalable small cell cultures, leading to the regeneration of full plants after the desired phenotypes have been optimized.

## Materials and methods

### Plant materials and growth condition

*Nicotiana benthamiana* and Maize B73 were grown in a Conviron growth chamber (Conviron, Winnipeg, Canada) at the University of Illinois with a 16-hour light and 8-hour dark photoperiod. The temperatures during the light and dark periods were 26 °C and 22 °C, respectively. The relative humidity was consistently maintained at 50% and the light intensity provided was 100 µmol m^−2^ sec^−1^, measured as photosynthetic photon flux density.

*N. benthamiana* plants used for callus generation in this study were germinated on 1⁄2 Murashige & Skoog (MS) media plates (Murashige and Skoog 1962) containing 2% sucrose under 16 h/8 h, 22 °C /18 ^°^C, light/dark conditions with 100 μmol m^−2^ s^−1^ light intensity. Leaf explants (0.5 x 0.5) cm were excised from approximately 2-week-old plants using aseptic technique. Explants were subsequently placed on MS plates containing 30 g/L sucrose, 0.1 g/L myo-inositol, 0.18 g/L KH_2_PO_4_, 1 mg/L Thiamine, 0.11 mg/L 2,4 D, pH 5.8 to induce callus generation. The plates were kept under continuous light at 25 °C for 2 weeks, and the resulting calli were transferred and maintained under continuous light at 25 ^°^C with a shaking speed of 120 rpm (An 1985).

### Plasmids used in this study

A0502: pMOD_A0502 (#91012, Addgene) for CRISPR knockout system in *N. benthamiana*, B2103: pMOD_B2013 (#91061, Addgene) for CRISPR knockout and activation systems in plants, C0000: pMOD_C0000 (#91081, Addgene) for CRISPR knockout and activation systems in plants, D100: pTRANS_100 (#91198, Addgene) for protoplast system, T230: pTRANS_230 (obtained from Dr. Voytas’s lab (Čermák et al. 2017)) for callus system, A1510: pMOD_A1510 (#91036, Addgene) CRISPR knockout system in maize, A3701: pMOD_A3701 (#91052, Addgene) for CRISPR activation system in *N. benthamiana*, A4110: pMOD_A4110 (#91056, Addgene) CRISPR activation system in maize. The detailed of the above plasmids can be found in the previous study (Čermák et al. 2017) and in Supplementary Table S1. pMDC43 and pMDC43-OWD can be found in the previous publications (Zhai et al. 2017b, 2021) for use in the transformation of *N. benthamiana* callus suspension cells, and the details of the sequences can be found in the Supplementary materials. Binary vector pPTN1586 was constructed by GoldenBraid modular assembly (Sarrion-Perdigones et al. 2013). Each gene of interest was synthesized by GenScript Biotech (Piscataway, NJ) to be GoldenBraid-domesticated and codon-optimized for sorghum. Genes used in pPTN1586 are Cuphea avigera var. pulcherrima DGAT1 (CpuDGAT1, ANN46862.1), sorghum Wrinkled1 (SbWRI1, XP_002450194.1), sesame oleosin (SiOle, Q9XHP2.1), Cuphea palustris thioesterase (Thio14, AAC49180.1), and Cuphea avigera var. pulcherrima class B lysophosphatidic acid acyltransferase (CpuLPATB, ALM22873.1). One amino acid residue in SbWRI1 was changed (K10R) for protein stability (Zhai et al. 2017a). pZP212 (pPTN1586C) served as an empty vector used as a control plasmid during the transfection of maize protoplast cells with pPTN1586. Maps for pPTN1586 and pZP212 (pPTN1586C) are provided in the Supplementary materials. pZP212 (Hajdukiewicz et al. 1994) (pPTN1586C) serves as an empty vector used as a control plasmid during the transfection of maize protoplast cells with pPTN1586. Detailed sequences of pPTN1586 and pZP212 can be found in the Supplementary materials.

### Automation methods and instruments

iBioFAB workflows were encoded into *Momentum* software (Thermo Scientific™, Waltham, MA, USA) (Enghiad et al. 2022), which coordinates instruments, controls the Thermo Fisher F5 robotic arm, and manages plate movement and tracking. The Beckman Coulter Echo 550 instrument (Beckman Coulter, Brea, CA, USA) was used for DNA cloning applications while all manipulations of plant cell cultures were performed on the Tecan Fluent 1080 robotic liquid handler (Tecan, Männedorf, Switzerland). The Tecan Fluent 1080 is equipped with a Pickolo (SciRobotics, Israel) colony picker.

Incubation steps were carried out in a Thermo Fisher Cytomat 2C automated incubator and centrifugation was performed in an Agilent Microplate Centrifuge (Agilent, Santa Clara, CA, USA). Growth of *Agrobacterium* cell cultures was quantified using a Tecan Infinite plate reader. All instruments are integrated onto a single platform and movement between instruments was performed by the Thermo Fisher F5 robotic arm. A 3D model of the integrated iBioFAB platform and flowchart of all automated workflows can be found in Supplementary Figures S1 and S2.

### Gene cloning, genomic DNA extraction and chlorophyll fluorescence measurement

Benchling software was used to generate CRISPR guide RNAs (gRNAs) for the *WRI1* gene (Maize: *Zm00001d037760; N. benthamiana: NbS00061229g0004*) and the *DGAT1* gene (Maize: *Zm00001d005016; N. benthamiana:* NbS00004767g0010. Additionally, CRISPR-P v2.0 (Lei et al. 2014) was employed to generate two gRNAs targeting *HCF136* gene (*N. benthamiana: NbS00049766g0015*). Plasmids containing gRNAs were cloned using the previously reported protocol (Čermák et al. 2017) and the detailed of protocol can be found in Supplementary Method 1, which was translated into an automation workflow (Supplementary Figure S1A). For assembly of B plasmid, a Beckman Coulter Echo 550 acoustic liquid handler was used to mix gRNA cassette PCR reactions, plasmid B and cloning reactions, then a Tecan Fluent was used to transform assembled plasmid B into *E. coli.* Next, plasmid B with two gRNA sequences was extracted from an overnight culture of *E. coli* using the Tecan Fluent, following our previously reported automated protocol (Enghiad et al. 2022). Sanger sequencing was employed to confirm B plasmids before subsequent assembly. A second round Golden Gate reaction was employed to assemble plasmid modules A, B with gRNAs, and C into the pTRANS backbone using the Echo 550 instrument again followed by transformation into *E. coli* using the Tecan Fluent. In summary, 15 vectors were cloned using FAST-PB pipeline are listed in Supplementary Table S1 and gRNAs and primers are listed in Supplementary Table S2.

Genomic DNA was isolated from plant cells using the Promega Wizard® SV 96 Genomic DNA Purification System (Promega Corporation, Madison, WI, USA) following the manufacturer’s protocol using the Tecan Fluent (Supplementary Figure S2E).

Subsequently, Tecan Fluent Flexible-Channel Arm (FCA) arm transferred 5 uL of the DNA extracts into a new 96-well plate for DNA measurements using the high-throughput Lunatic UV/Vis absorbance spectrometer Microfluidic system (Unchained Labs, Pleasan).

After transfection for 24 hours with *HCF136* in maize or *N. benthamiana*, we utilized the Tecan Infinite Plate reader on the iBioFAB platform to measure chlorophyll fluorescence after overnight dark incubation of cell culture. This measurement was performed by setting the Excitation Wavelength to 650 nm and the Emission Wavelength to 675 nm. The output yielded two datasets: one representing the intensity of chlorophyll fluorescence and the other reflecting cell intensity, as indicated by A600 measurements. Subsequently, the final result value was calculated as the ratio of the intensity of chlorophyll fluorescence to cell intensity.

### Protoplast isolation and transfection

To isolate protoplasts from Maize B73, 30 leaves were collected from 14-day-old etiolated seedlings then sliced into pieces with 1 mm thickness and evenly distributed into a 24 square-well V-bottom plate. *N. benthamiana* protoplasts were isolated from five 8-week-old leaves cut into 2 mm thickness and distributed to another same type of plate. After slicing the leaves and distributing them into two 24-well plates (one for maize and the other for *N. benthamiana*), the remaining isolation steps were performed on the iBioFAB platform (Supplementary Figures S2C-D). First, the FCA arm of Tecan Fluent distributed 2 mL of enzyme solution into each well of a 24-well plate followed by shaking for four hours at 100 RPM at room temperature without light in the incubator of the Tecan Fluent. Next, the digested leaf solution was filtered on a 24-well AcroPrep filter plate with 30-40 µm pore volume (Pall Corporation, Port Washington, New York, USA) using the Te-VacS vacuum separation module integrated into the Tecan Fluent system. The filtered solution was then centrifuged on an Agilent automated microplate centrifuge (Agilent, Santa Clara, CA, USA) for 2 min at 150 rcf. The Tecan Fluent was again used to remove the supernatant and wash cells by adding 200 µL of W5 solution to each well. Following another round of centrifugation and supernatant removal, the cells were resuspended into 200 µL of W5 solution. All cells from all 24 wells were then combined into a single tube and centrifuged again. After supernatant removal, the cells were resuspended into 5 mL of MMG solution using the FCA arm, and the cell concentration was determined using a hemocytometer (Thermo Scientific, Waltham, MA, USA).

Next, the FCA arm slowly distributed 100 µL of isolated protoplasts into each well of a 96-well plate. To optimize transfection efficiency, three different average number molecular weights of PEG (2050, 3350, and 4000) were used: PEG 2050 (Sigma-Aldrich, Lot #: BCBW7040), PEG 3350 (Sigma-Aldrich, Lot #: MKCL5061), and PEG 4000 (Sigma-Aldrich, Lot #: BCCF2031). These three types of PEG and one type of plasmid p201GFP-Cas9 (Jacobs et al. 2015) were gently added into each well of 96-well plate containing protoplast cells and then slowly mixing them, according to a predefined worklist using the Momentum^TM^ software. The transformation mixtures were incubated in the dark at room temperature for 30 min. Subsequently, the FCA arm added 600 µL of W5 solution to each well and mixed gently to stop the transformation reaction. The F5 robotic arm moved the plate to the centrifuge followed by centrifugation for 2 min at 150 rcf (Relative Centrifugal Field) and then removal of supernatants and resuspension into fresh W5 solution. After second wash, plates were centrifuged again for 2 min at 150 rcf and cells were resuspended into 100 µL of W5 solution through using the Tecan Fluent system. After determining that PEG 2050 yielded the highest transfection efficiency, we used this optimal PEG type to transfect cells in both maize B73 and *N. benthamiana*. All automated transfection procedures with same PEG type 2050 but different plasmids were conducted as previously described. In maize B73, we employed eight types of plasmids (A4110, A4110-DGAT1, A4110-WRI1, pPTN1586C, pPTN1586, A1510, A1510-HCF136, p201GFP-Cas9 (Jacobs et al. 2015)), with each type having four to ten replicates. Similarly, *for N. benthamiana* transfection, we used six types of plasmids (P-A3701, P-A3701-DGAT1, P-A3701-WRI1, A0502, A0502-HCF136, p201GFP-Cas9), also with four to ten replicates for each type.

### *N. benthamiana* callus cell culture transformation

Two seven-week-old calli were placed in a single well of a 6-well plate, resulting in two calli per well. Tecan Fluent was used to add 3 mL MS liquid media to each well. The plate was then exposed to continuous light for a period of 20 days while shaking at 200 rpm (Revolutions Per Minute). Prior to transformation, the Pickolo colony picker (SciRobotics, Kfar Saba, HaMerkaz, Israel) was used to pick *Agrobacterium tumefaciens* colonies from an agar plate containing rifampicin and kanamycin antibiotics into a 96-deepwell plate containing 1 mL LB media per well. Specifically, six colonies were selected from six different plates, namely A3701, A3701-WRI1, A3701-DGAT1, A1510-HCF136, pMDC43, and pMDC43-OWD. The F5 robotic arm transferred the plate to the Thermo Scientific™ Cytomat™_6K automated incubator (Thermo Scientific, Waltham, MA, USA), where it underwent overnight outgrowth at 200 rpm. The following day, the 96-deep well plate was taken out from the incubator and placed on the dock of the Tecan Fluent. Subsequently, the FCA arm of the Tecan Fluent transferred 100 μl of the overnight culture to a new 96-well plate for optical density (OD) measurement at 600 nm using Tecan infinite plate reader (Männedorf, Switzerland) on the iBioFAB platform. Following the OD measurement, if the OD of the cell culture was within the range of 0.4-1, the optimal overnight culture was introduced to the ten-week-old callus suspension cells in the 6-well plate. The co-culture was conducted under light conditions for two days. Next, co-culture cells were washed four times with MS liquid media using the Tecan Fluent. After washing, calli transformed with A3701, A3701-WRI1, A3701-DGAT1, and A1510-HCF136 were placed on 2 mg/L phosphinothricin (PPT) selection MS media plates, while those transformed with pMDC43 and pMDC43-OWD were placed on a 15 mg/L hygromycin-selected MS media plates. The automated procedures are demonstrated in Supplementary Figure S2B. Shoot and root regeneration procedures were followed in the previous study (Clemente 2006). Briefly, genetically engineered calli were transferred into the shoot media (MS Salts & MS vitamins + 30 g sucrose + 2 mg/L kinetin + 1 mg/L IAA+ 400 mg/L Timentin + 2.0 mg/L glufosinate ammonium-for bar gene selection- or 20 mg/L hygromycin selection). After 10-16 weeks, shoots were generated then transferred to the root media (MS salts & vitamins + 30g sucrose+ 200 mg/L timentin + 2.0 mg/L glufosinate ammonium- for bar gene selection- or 20 mg/L hygromycin selection- for rooting). After 5-8 weeks, fully rooted plants were transferred to pots with PRO-MIX BX BIOFUNGICIDE MYCORRHIZAE soil in the growth chamber.

### RNA extraction and RT-qPCR analysis

RNA was extracted from callus and protoplast cultures using the ZR Plant RNA Miniprep™ kit (R2024, Zymo Research, CA, USA) and TRIzol™ Reagent (15596026, Thermo Scientific, Waltham, Massachusetts, USA), respectively. The single-stranded cDNA was synthesized using the High-Capacity cDNA Reverse Transcription Kit (ThermoFisher Scientific; Waltham, Massachusetts, USA). Real-time PCR (qPCR) was performed using the Power SYBR® Green PCR Master Mix (ThermoFisher Scientific) in a LightCycler 480 instrument (Roche; Indianapolis, IN, USA). Gene expression levels were normalized to the expression of the constitutively expressed reference genes (Table S2). Relative gene expression was calculated following previously published methods (Livak and Schmittgen 2001; Dong et al. 2020). The qPCR primers are shown in Supplementary Table S2.

### Lipid extraction and LC-MS/MS analysis

Lipid extraction from plant tissues and protoplasts were performed as described elsewhere (Zhai et al. 2018). Briefly, 700 µL of extraction solvent (chloroform: methanol: formic acid 2:1:0.1 v/v/v) and 350 µL of 1M KCl with 0.2M H_3_PO_4_ were added for liquid-liquid partition. This was vortexed for 30 min, centrifuged at 20,000 x g for 10 min, then the lower layer was taken and dried using a SpeedVac vacuum concentrator (Thermo Scientific, USA). The samples were redissolved in 200 µL of an LC-MS grade solvent mixture of isopropanol/acetonitrile/water 65/30/5, added to an HPCL vial insert, and 2 µL SPLASH^TM^ LIPIDOMIX® Mass Spec Standard (Avanti Polar Lipids was added to each of the samples for an internal calibrant.

LC-MS/MS analysis was performed on a Vanquish^TM^ UHPLC coupled with Q Exactive^TM^ Orbitrap Mass Spectrometer (Thermo Scientific, USA) with HESI source. An Acquity UPLC^®^ BEH C18 column (2.1 × 100 mm, 1.7 μm) was kept at 45 °C. Solvent A was 60:40 (v/v) acetonitrile:water with 10 mM ammonium formate and 0.1 % formic acid, and solvent B was 90:10 (v/v) isopropanol:acetonitrile with 10 mM ammonium formate and 0.1 % formic acid. The gradient started with 15% phase B and increased to 50% at 2 min, then to 98% at 15.5 min. The column was washed at 98% phase B for 2 min, and continued with equilibration using 15% B from 17.6 to 20 min. Flow rate was kept at 250 μl/min.

For mass spectrometry analysis, the capillary temperature was set at 300 °C, for both positive and negative modes. Sheath gas flow rate was set to 35 and aux gas to 10. For positive scan, the spray voltage was 3.5 kV and for negative was 2.8 kV. Positive and negative data were collected by separate injections. Data was acquired by full MS scan followed by data dependent scans with fragmentation energy. Full MS scan range was *m/z* 150-1500. The AGC (Automatic Gain Control) target was set to 3e6. For data dependent MS^2^, the top 10 ions were selected for fragmentation at stepped NCE (Normalized collision energy) of 15, 25 and 35. The isolation window was *m/z* 1.0, resolution was set to 17,500, the AGC target was at 1e^5^, and dynamic exclusion was set as 5.0 s for triggered ions. Centroid mode was used for all data collection. Peak detection, alignment, and identification were performed on MS-Dial (ver 4.90) with built-in *in silico* LC/MS/MS based lipidomics database (Tsugawa et al. 2020). Identification was based on MS2 match, and the score cut off was set at 80%.

### GC-MS analysis

Qualitative, targeted fatty acids analysis was performed using an Agilent 6890N gas chromatography attached to a 5975B MS in the Metabolomics Laboratory of Roy J. Carver Biotechnology Center, University of Illinois at Urbana-Champaign, as previously described (Xue et al. 2020). One microliter injection of the sample was made into the column in a Pulsed Splitless mode, with the front inlet pressure elevated to 40 psi for 18 s. Helium was the carrier gas used. The front inlet, MS transfer line, MS source, and MS quad were maintained at 300 °C, 230 °C, 230 °C and 150 °C, respectively. The GC oven temperature protocol was as follows: 50 °C for 2 min, ramp up at 30 °C/min for a 2 min hold at 120 °C, a second ramp up at 30 °C/min for a 2 min hold at 180 °C, and a final ramp up at 30 °C/min for a 9.33 min hold at 250°C.

### Matrix application and high-throughput MALDI MS analysis

A matrix solution containing 45 mg/mL 2,5-dihydroxybenzoic acid dissolved in 70% methanol was deposited onto ITO-coated microscopy slides using an HTX M5-Sprayer (HTX Technologies). The sprayer temperature was set to 70 °C, with a flow rate of 0.1 ml min^-1^, track spacing of 3 mm, pressure of 10 psi, and a gas flow rate of 3 l min^-1^. One pass of the matrix was applied to the slides, with a final matrix density of 3.195 mg/mm^2^.

MALDI MS analysis was performed with a SolariX XR 7T FTICR mass spectrometer (mass spectrometer equipped with an APOLLO II dual MALDI/ESI source (Bruker). Data were acquired in positive-mode with a mass range of *m/z* 54-1,600, yielding a transient length of 0.524 s using a Smartbeam-II UV laser (355 nm) set to “Small” mode, generating a ∼100-µm diameter laser footprint. Each MALDI acquisition consisted of 500 laser shots at a frequency of 1,000 Hz. microMS was used as previously described to generate instrument stage coordinates and geometry files for all MALDI acquisitions of selected protoplasts with a distance filter of 200 µm (removed protoplasts located closer than 200 µm to each other from the target list) (Comi et al. 2017).

Peak picking and peak export were performed using Compass DataAnalysis 4.4 (Bruker) with a signal-to-noise ratio or 5 and a relative intensity threshold of 0.01%. Mass spectral binning was performed in custom Matlab scripts with a semicontinuous bin width of 3 ppm. Features were mass matched to a bulk LC-MS/MS database using a 5 ppm filter, and cells with fewer than 5 matched lipids were removed from the sample set.

### Statistical methods in this study

All the P-values provided were generated using the two-tailed Welch’s t-test. Statistical analysis, including volcano plot and heatmap analysis, was conducted using the web tool available at https://www.metaboanalyst.ca/ (Lu et al. 2023).

## Author contributions

All authors designed experiments, analyzed data, and assisted in the writing and editorial process. J.D., S.W.C., S.L., D.C.C., J.B., S.Z., K.P. performed the experiments. J.S., H.Z., J.V.S., and M.E.H. conceived and supervised the overall project.

## Acknowledgments

We express our gratitude to Benjamin Haas for generously sharing the gene cloning protocol and reference primers for RT-qPCR, which greatly contributed to the success of this study. We sincerely thank Michael John Root for sharing the plant regeneration protocol and technique support, which greatly contributed to *N. benthamiana* regeneration. We also extend our thanks to Dr. Thomas Clemente for providing the necessary information on plasmids (pPTN1586C and pPTN1586), which were instrumental in advancing our research. We would like to acknowledge Dr. Zhiyang Zhan and Dr. Yingqi Cai for providing the plasmids (pMDC43 and pMDC43-OWD).

## Supplementary information

### Materials

#### pMDC43 plasmid (referred to as 43)

Map of a control vector without any genes designed to alter plant metabolism. The details of the sequence can be found in the following link: https://benchling.com/s/seq-ynhIYe0uQCFpD6xAAuj4?m=slm-2lWxgjrKdKwnzn3R7ybC

#### pMDC43-OWD plasmid (referred to as 43-OWD)

Contains three genes targeted at plant metabolism, which are Oleosin, WRI1 and DGAT1. The details of the sequence can be found in the following link: https://benchling.com/s/seq-CNuD3g9jJb1rnoycCcGq?m=slm-nF6krntKZMu41ywKaHKX

#### pPTN1586

contains *DGAT1*, *WRI1*, *Oleosin, Thio14* and *LPATB* sequences. The schematic view of T-DNA for pPTN1586 is provided below. LB; left border, RB; right border, 35S; cauliflower mosaic virus 35S promoter, UBI4; sugarcane ubiquitin4 promoter, UBI1; maize ubiquitin1 promoter WR1; synthetic Wrinkled1-responsive promoter1 generated by10x repeated AW-box motifs with minimal 35S promoter, WR2; synthetic Wrinkled1-responsive promoter2 generated by10x repeated reverse AW-box motifs with minimal 35S promoter, 35Ster; cauliflower mosaic virus 35S terminator, OCSter; *Agrobacterium* Ocs terminator, NOSter; *Agrobacterium* nopaline synthase (Nos) terminator, tSbACT; sorghum actin1 terminator. The details of the sequence can be found in the following link: https://benchling.com/s/seq-He5x2ty62Gw3L3KYGlIv?m=slm-zpOzF773ZPYpFiGDgBhy

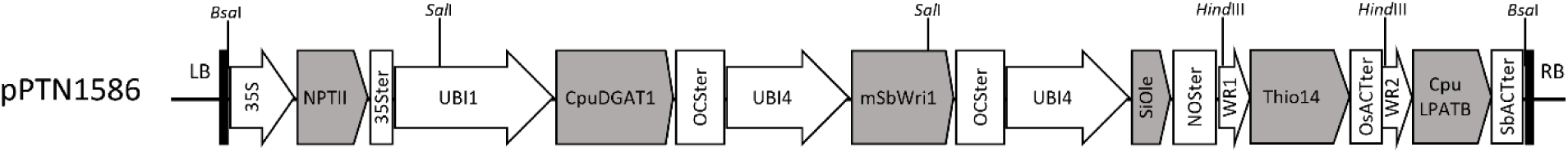

#### pZP212 plasmid

A control vector without any genes designed to alter plant metabolism. The details of the sequence can be found in the following link: https://benchling.com/s/seq-vRo728AuHezhUaZo28RD?m=slm-j6HekX4GC6mh6ULw5KoQ

## Methods

### Supplementary method 1: Golden Gate Cloning

The workflow for generating CRISPR-constructs is based on the Golden Gate cloning method described by Cermak et al.(Čermák et al. 2017). The details of the cloning method can be found in the provided link below. Please note that in this study, we did not perform the ’Assembly of Module C vectors - Gene targeting’ as outlined in the following protocol.

https://benchling.com/s/prt-UDhXqCcWdlsnXiq2YRZH/edit

### Supplementary method 2: Protoplast isolation and transfection method

Protoplast isolation and transfection method can be found at this online link: http://patronlab.org/wp-content/uploads/2017/03/PREPARATION-AND-TRANSFECTION-OF.pdf

## Supplementary Figures

**Supplementary Figure S1.**
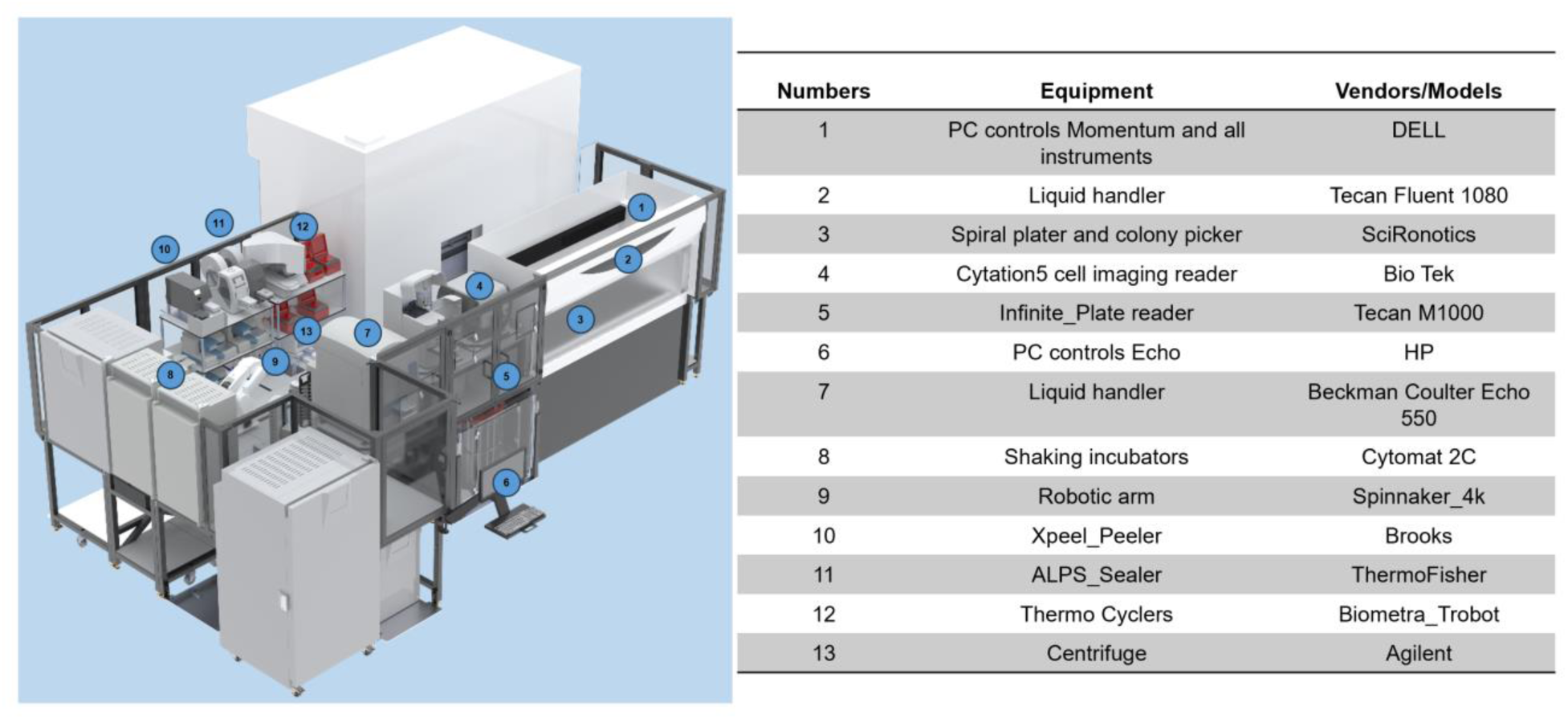
Instruments and software were utilized on the iBioFAB platform for this study. Layout of the integrated hardware (#1-13) inside the iBioFAB.

**Supplementary Figure S2.**
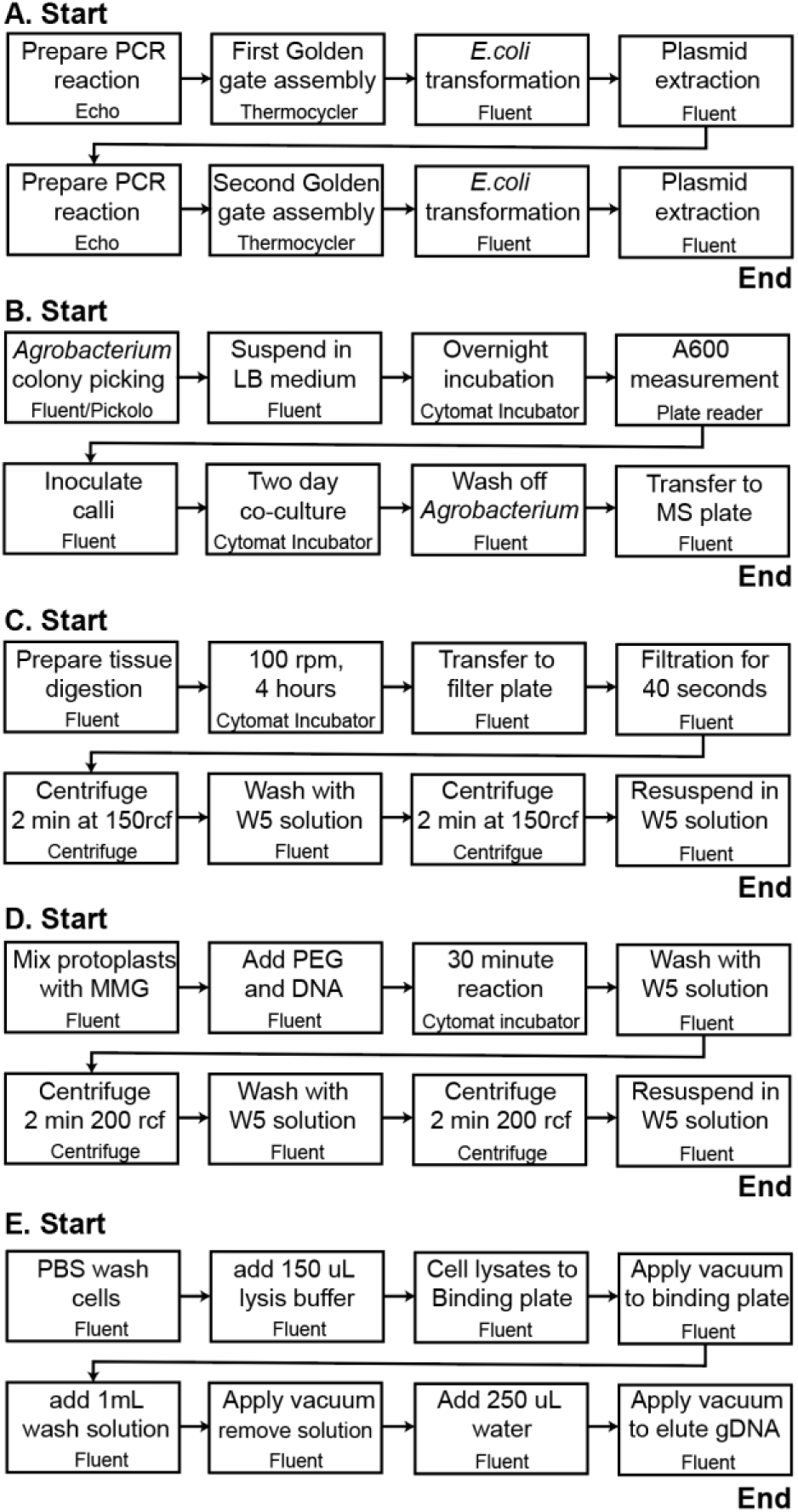
Detailed workflow of each automation modules. (A) Gene cloning through the Golden Gate cloning method. (B) Callus transformation. (C) Protoplast isolation from the etiolated maize B73 and *N. benthamiana*. (D) Protoplast transfection with plasmids. (E) Genomic DNA extraction. The top of each box describes the workflow step while the utilized equipment is shown in smaller font at the bottom of each box.

**Supplementary Figure S3.**
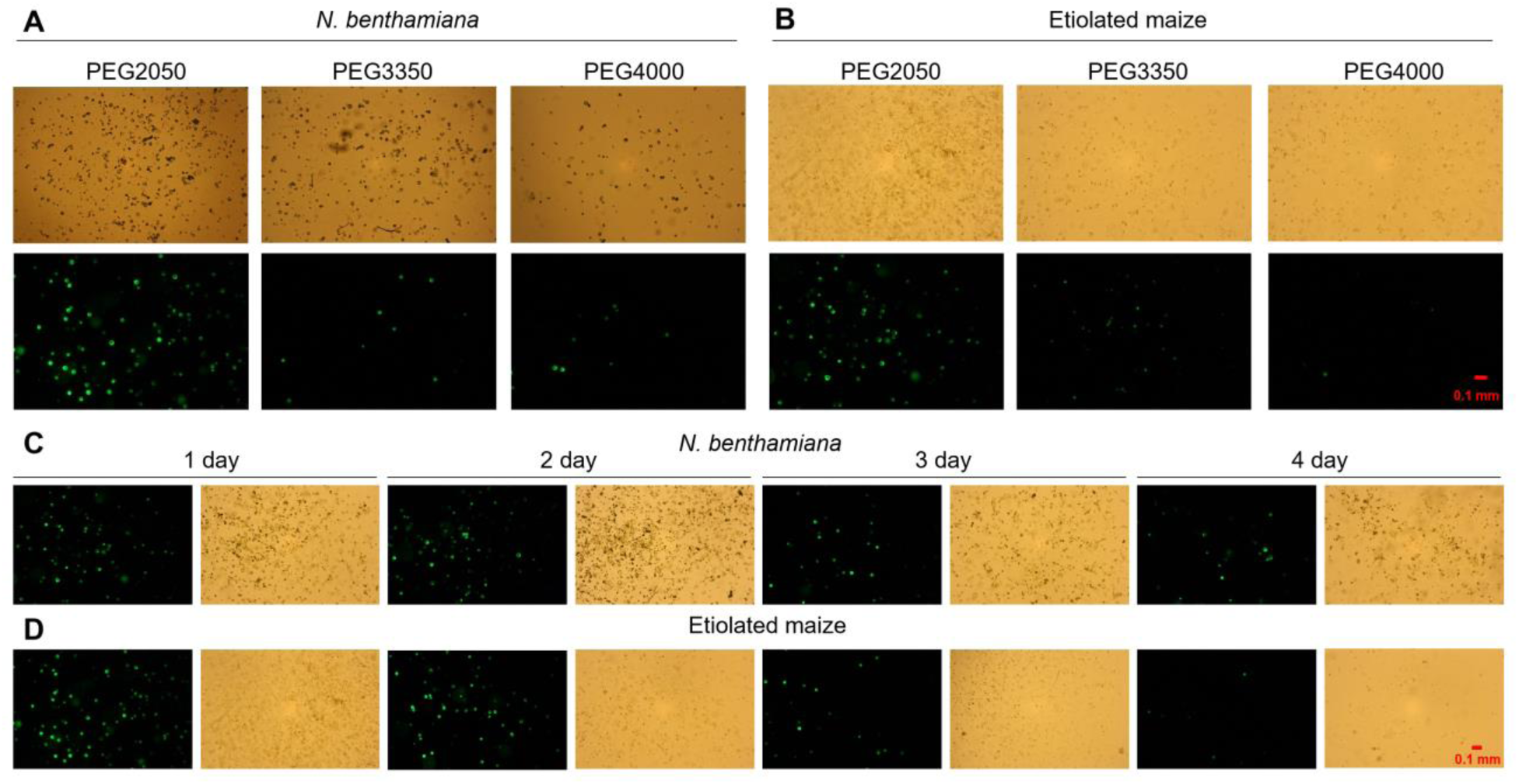
Transfection with PEG treatment and GFP expression in both *N. benthamiana* and maize. (**A-B**) Transfection with three types of PEG (2050, 3350 and 4000) in *N. benthamiana* (**A**) and maize (**B**). Images at the top were captured under bright-field illumination, whereas the images in the second row were acquired using the GFP channel. (**C-D**) GFP expression after transfection with Cas9 vector (p201G Cas9) from first day to fourth day in *N. benthamiana* (**C**) and maize (**D**). Each day has two images, left is a GFP picture and right is a bright field image.

**Supplementary Figure S4.**
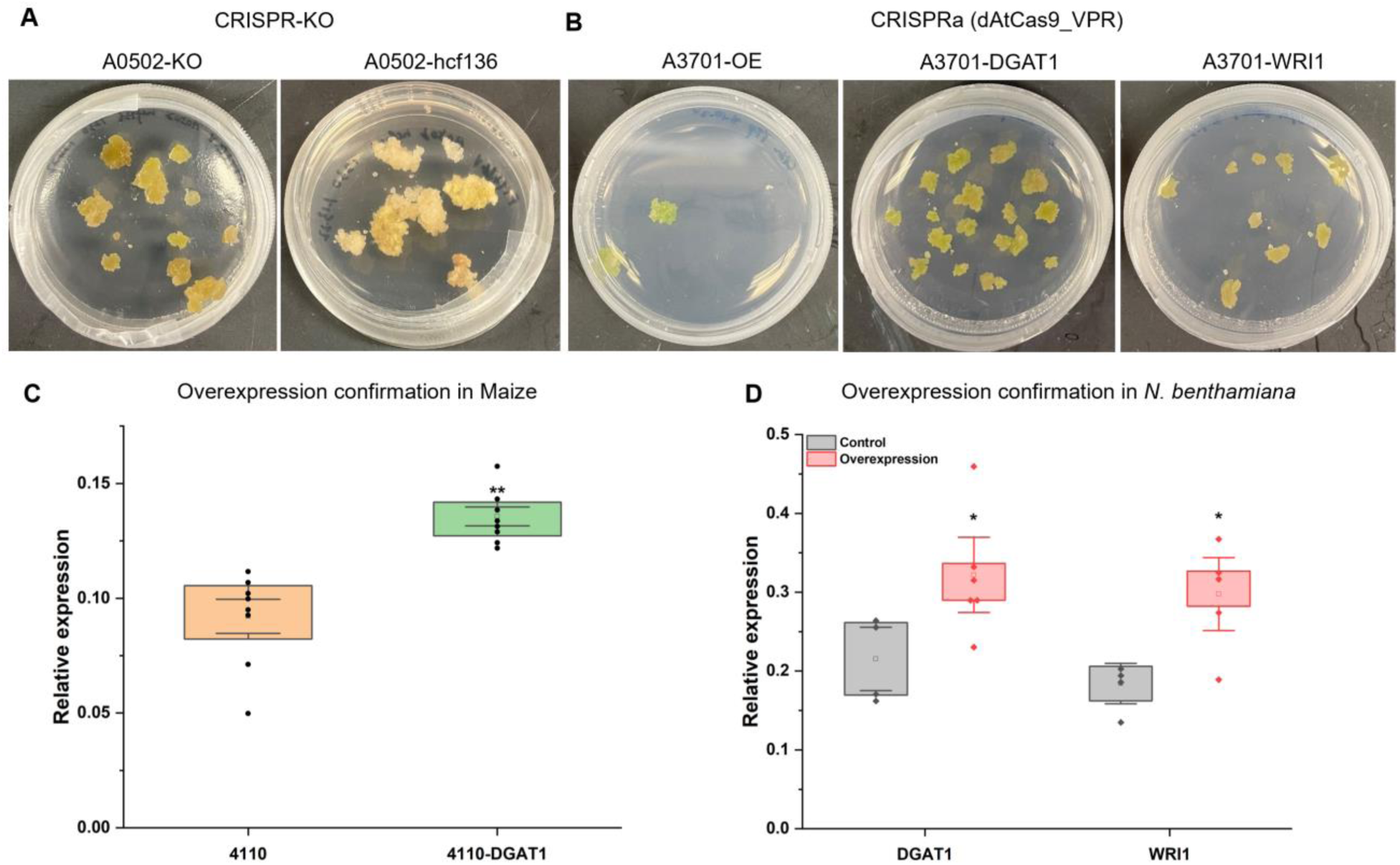
Genetically engineered calli generated on the iBioFAB platform. (**A**) Calli harboring the knockout gene *HCF136*, generated with the CRISPR knockout system, thrive on MS media supplemented with phosphinothricin (PPT). (**B**) Calli with the overexpression of lipid genes, induced by the CRISPR activation system, grow on MS media supplemented with PPT. (**C**) RT-qPCR results show the overexpression of *DGAT1* gene achieved using CRISPR activation in maize protoplast cells. Biological replicate n = 8. (**D**) RT-qPCR results demonstrate the successful overexpression of *WRI1* or *DGAT1* gene through CRISPR activation in *N. benthamiana*. Biological replicate n = 4. Error bars represent standard error. Asterisks indicate: \*\*\**P* < 0.001; \*\**P* < 0.01; \**P* < 0.05, calculated using two-tailed Welch’s t-test.

**Supplementary Figure S5.**
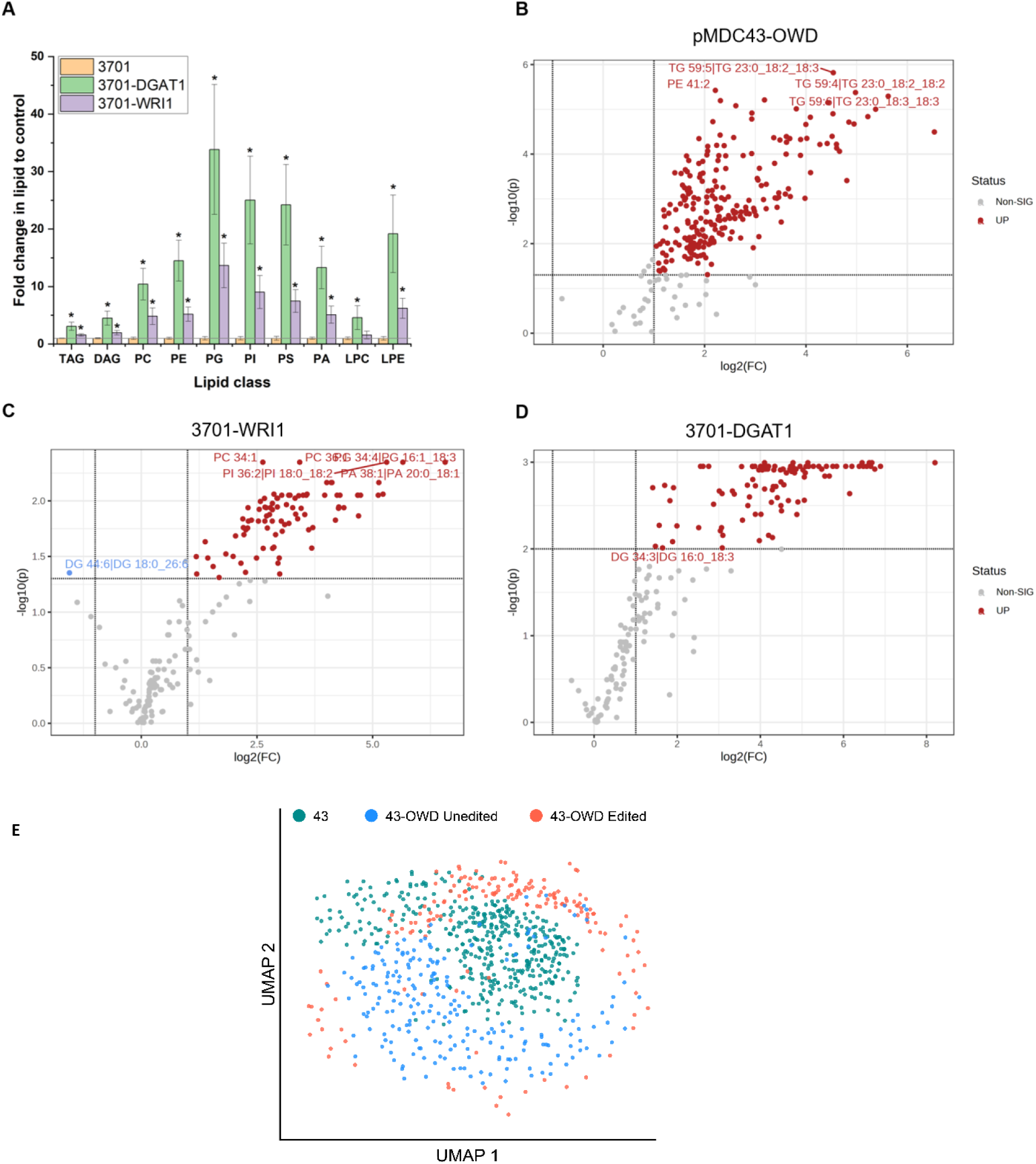
Lipid quantification and statistical analysis. (**A**) Lipid quantification via LC-MS/MS of genetically engineered calli protoplast cells: control 3701, *DGAT1* overexpression 3701-DGAT, and *WRI1* overexpression 3701-WRI1. Volcano plots generated from statistical analysis of (**B**) OWD (*Oleosin*, *WRI1*, and *DGAT1*) overexpression using the vector pMDC43-OWD. (**C**) *WRI1* overexpression, and (**D**) *DGAT1* overexpression. Biological replicates n = 6. In this context, FC represents fold change, and p signifies significance (*P* < 0.05). Error bars represent standard error. Asterisks indicate: \*\*\**P* < 0.001; \*\**P* < 0.01; \**P* < 0.05, calculated using two-tailed Welch’s t-test. (**E**) Uniform Manifold Approximation and Projection (UMAP) visualization illustrating the distribution of cells transfected with the control vector pMDC43 (43), three-gene stack pMDC43-OWD cells that remained unedited (43-OWD unedited), and pMDC43-OWD cells that were successfully edited (43-OWD edited). Each point represents an individual cell. The pMDC43-OWD dataset was subdivided into “unedited” and “edited” by comparing each individual protoplast’s lipid profiles to the mean of the pMDC43 control group.

---

**Supplementary Table S1.** List of plasmids used in this study.

| Cell systems | *N. benthamiana* | Maize |
| --- | --- | --- |
| **CRISPR knockout system** |  |  |
| Protoplast system | A0502-B2103-C0000-D100<br>(A0502) | A1510-B2103-C0000-D100<br>(A1510) |
|  | A0502-B2103-HCF136-C0000-D100<br>(A0502-HCF136) | A1510-B2103-HCF136-C0000-D100<br>(A1510-HCF136) |
| Callus system | A0502-B2103-C0000-T230<br>(A0502) | N/A |
|  | A0502-B2103-HCF136-C0000-T230<br>(A0502-HCF136) | N/A |
| **CRISPR activation system** |  |  |
| Protoplast system | A3701-B2103-C0000-D100<br>(A3701) | A4110-B2103-C0000-D100<br>(A4110) |
|  | A3701-B2103-DGAT1-C0000-D100<br>(A3701-DGAT1) | A4110-B2103-DGAT1-C0000-D100<br>(A4110-DGAT1) |
|  | A3701-B2103-WRI1-C0000-D100<br>(A3701-WRI1) | A4110-B2103-WRI1-C0000-D100<br>(A4110-WRI1) |
| Callus system | A3701-B2103-C0000-T230<br>(A3701) | N/A |
|  | A3701-B2103-DGAT1-C0000-T230<br>(A3701-DGAT1) | N/A |
|  | A3701-B2103-WRI1-C0000-T230<br>(A3701-WRI1) | N/A |

Note: The gene name enclosed in brackets represents the short name of the full assembly of several plasmids, and "N/A" is used to indicate that there is no assembly of plasmids for the gene in that cell system.

**Supplementary Table S2.**
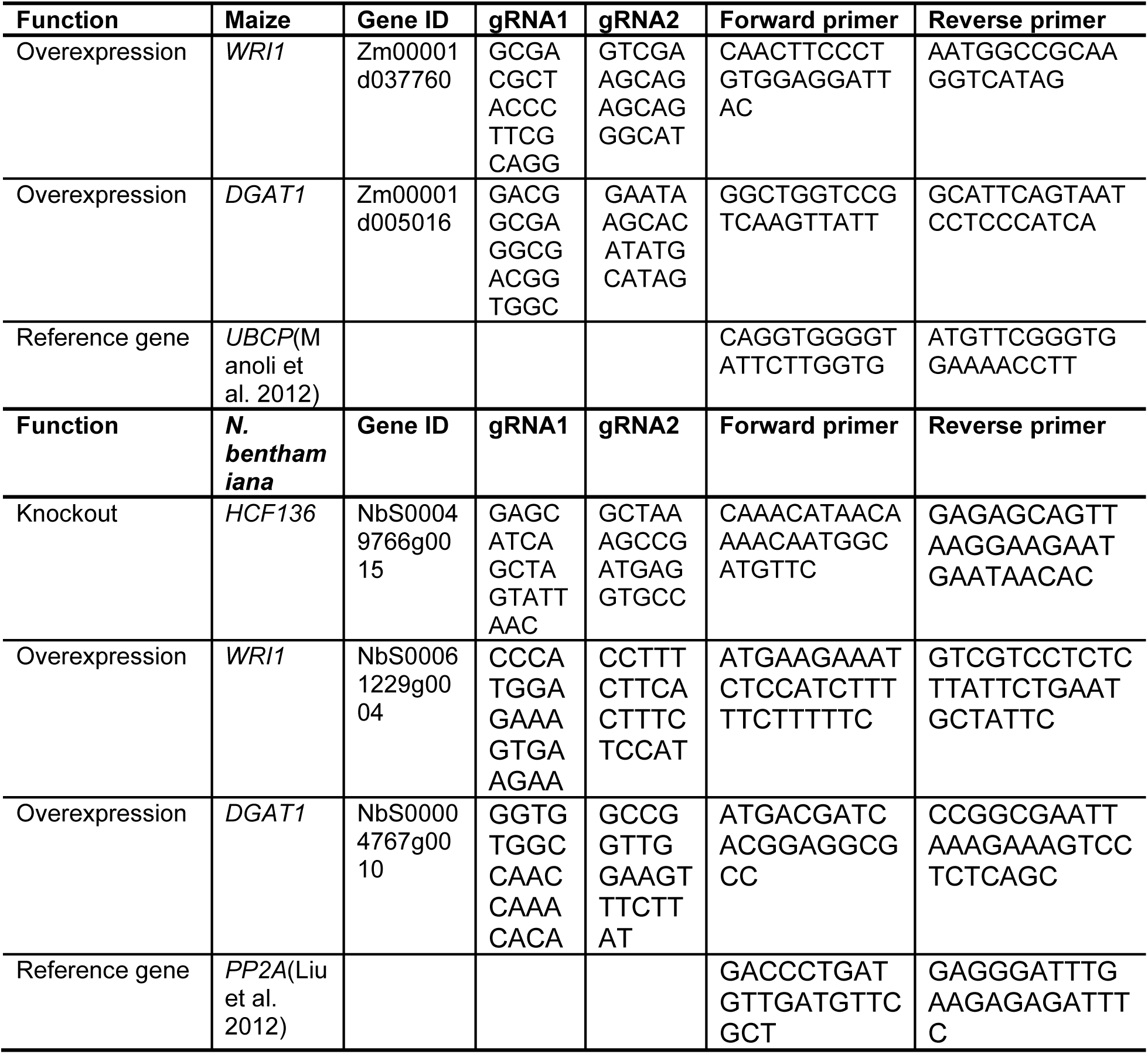
List of gRNAs and primers used in this study.

**Supplementary Table S3.**
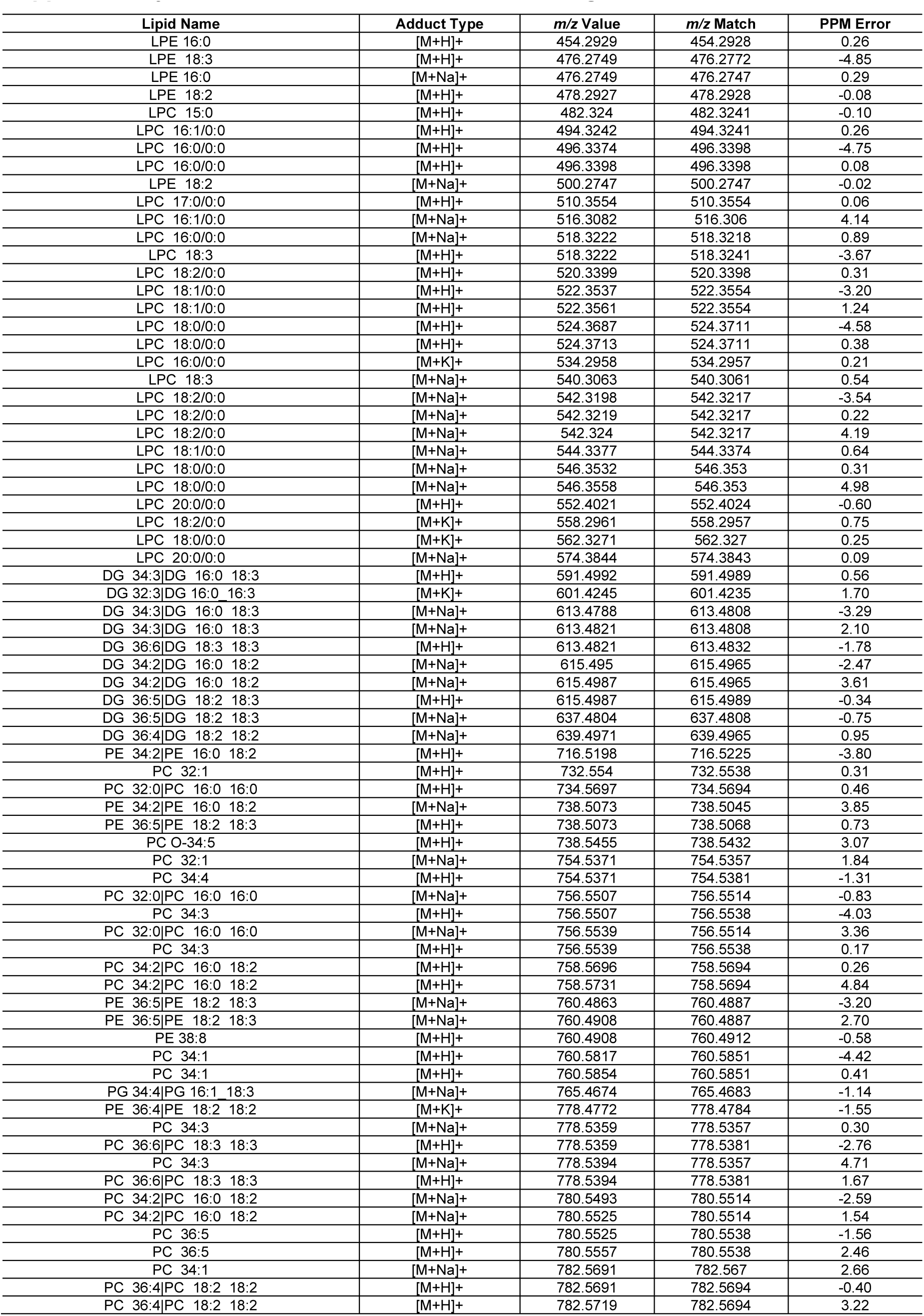
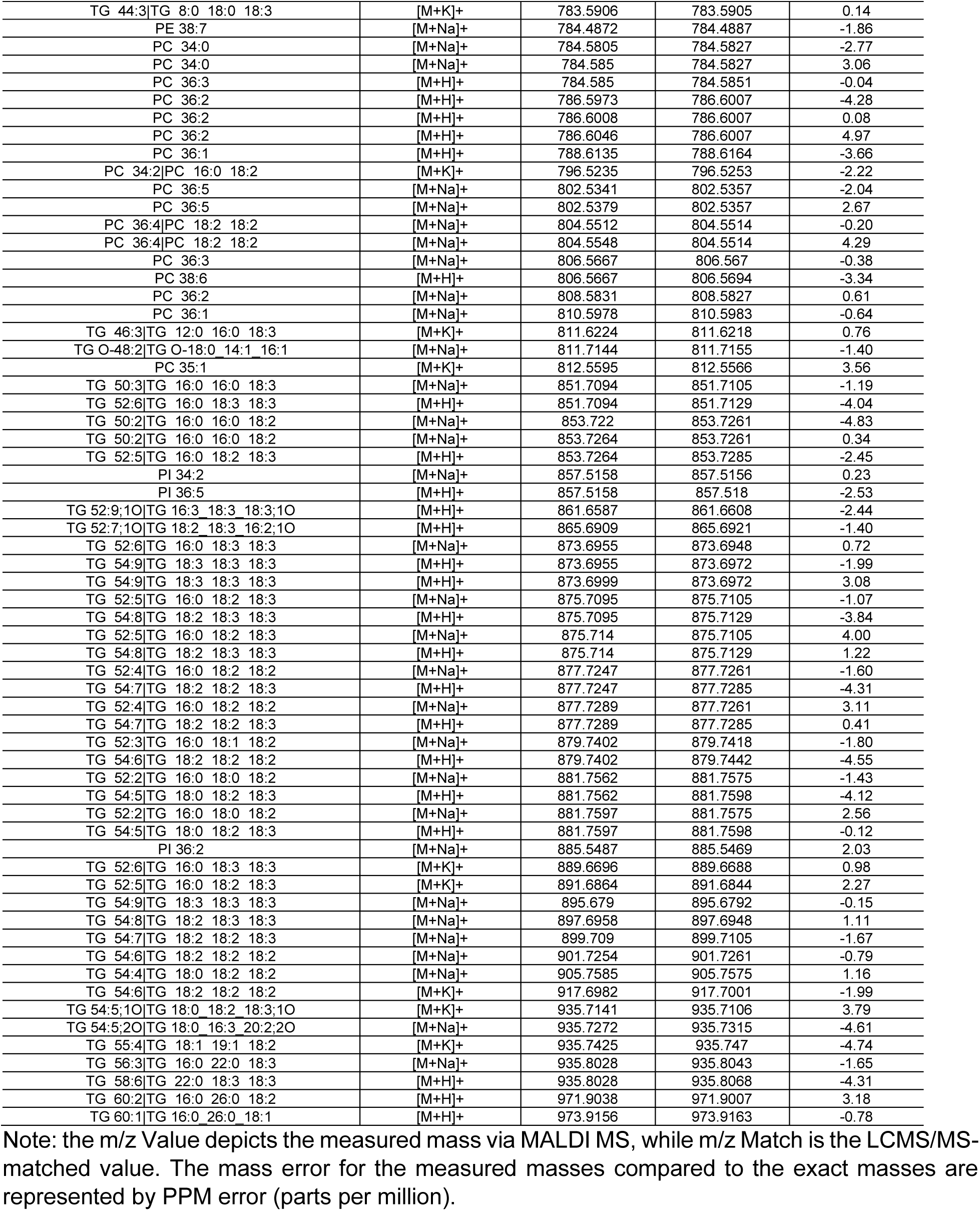
MALDI MS mass match assignments.

| Lipid Name | Adduct Type | m/z Value | m/z Match | PPM Error |
| --- | --- | --- | --- | --- |
| LPE 16:0 | [M+H]<sup>+</sup> | 454.2929 | 454.2928 | 0.26 |
| LPE 18:3 | [M+H]<sup>+</sup> | 476.2749 | 476.2772 | -4.85 |
| LPE 16:0 | [M+Na]<sup>+</sup> | 476.2749 | 476.2747 | 0.29 |
| LPE 18:2 | [M+H]<sup>+</sup> | 478.2927 | 478.2928 | -0.08 |
| LPC 15:0 | [M+H]<sup>+</sup> | 482.324 | 482.3241 | -0.10 |
| LPC 16:1/0:0 | [M+H]<sup>+</sup> | 494.3242 | 494.3241 | 0.26 |
| LPC 16:0/0:0 | [M+H]<sup>+</sup> | 496.3374 | 496.3398 | -4.75 |
| LPC 16:0/0:0 | [M+H]<sup>+</sup> | 496.3398 | 496.3398 | 0.08 |
| LPE 18:2 | [M+Na]<sup>+</sup> | 500.2747 | 500.2747 | -0.02 |
| LPC 17:0/0:0 | [M+H]<sup>+</sup> | 510.3554 | 510.3554 | 0.06 |
| LPC 16:1/0:0 | [M+Na]<sup>+</sup> | 516.3082 | 516.306 | 4.14 |
| LPC 16:0/0:0 | [M+Na]<sup>+</sup> | 518.3222 | 518.3218 | 0.89 |
| LPC 18:3 | [M+H]<sup>+</sup> | 518.3222 | 518.3241 | -3.67 |
| LPC 18:2/0:0 | [M+H]<sup>+</sup> | 520.3399 | 520.3398 | 0.31 |
| LPC 18:1/0:0 | [M+H]<sup>+</sup> | 522.3537 | 522.3554 | -3.20 |
| LPC 18:1/0:0 | [M+H]<sup>+</sup> | 522.3561 | 522.3554 | 1.24 |
| LPC 18:0/0:0 | [M+H]<sup>+</sup> | 524.3687 | 524.3711 | -4.58 |
| LPC 18:0/0:0 | [M+H]<sup>+</sup> | 524.3713 | 524.3711 | 0.38 |
| LPC 16:0/0:0 | [M+K]<sup>+</sup> | 534.2958 | 534.2957 | 0.21 |
| LPC 18:3 | [M+Na]<sup>+</sup> | 540.3063 | 540.3061 | 0.54 |
| LPC 18:2/0:0 | [M+Na]<sup>+</sup> | 542.3198 | 542.3217 | -3.54 |
| LPC 18:2/0:0 | [M+Na]<sup>+</sup> | 542.3219 | 542.3217 | 0.22 |
| LPC 18:2/0:0 | [M+Na]<sup>+</sup> | 542.324 | 542.3217 | 4.19 |
| LPC 18:1/0:0 | [M+Na]<sup>+</sup> | 544.3377 | 544.3374 | 0.64 |
| LPC 18:0/0:0 | [M+Na]<sup>+</sup> | 546.3532 | 546.353 | 0.31 |
| LPC 18:0/0:0 | [M+Na]<sup>+</sup> | 546.3558 | 546.353 | 4.98 |
| LPC 20:0/0:0 | [M+H]<sup>+</sup> | 552.4021 | 552.4024 | -0.60 |
| LPC 18:2/0:0 | [M+K]<sup>+</sup> | 558.2961 | 558.2957 | 0.75 |
| LPC 18:0/0:0 | [M+K]<sup>+</sup> | 562.3271 | 562.327 | 0.25 |
| LPC 20:0/0:0 | [M+Na]<sup>+</sup> | 574.3844 | 574.3843 | 0.09 |
| DG 34:3/DG 16:0 18:3 | [M+H]<sup>+</sup> | 591.4992 | 591.4989 | 0.56 |
| DG 32:3/DG 16:0 16:3 | [M+K]<sup>+</sup> | 601.4245 | 601.4235 | 1.70 |
| DG 34:3/DG 16:0 18:3 | [M+Na]<sup>+</sup> | 613.4788 | 613.4808 | -3.29 |
| DG 34:3/DG 16:0 18:3 | [M+Na]<sup>+</sup> | 613.4821 | 613.4808 | 2.10 |
| DG 36:6/DG 18:3 18:3 | [M+H]<sup>+</sup> | 613.4821 | 613.4832 | -1.78 |
| DG 34:2/DG 16:0 18:2 | [M+Na]<sup>+</sup> | 615.495 | 615.4965 | -2.47 |
| DG 34:2/DG 16:0 18:2 | [M+Na]<sup>+</sup> | 615.4987 | 615.4965 | 3.61 |
| DG 36:5/DG 18:2 18:3 | [M+H]<sup>+</sup> | 615.4987 | 615.4989 | -0.34 |
| DG 36:5/DG 18:2 18:3 | [M+Na]<sup>+</sup> | 637.4804 | 637.4808 | -0.75 |
| DG 36:4/DG 18:2 18:2 | [M+Na]<sup>+</sup> | 639.4971 | 639.4965 | 0.95 |
| PE 34:2/PE 16:0 18:2 | [M+H]<sup>+</sup> | 716.5198 | 716.5225 | -3.80 |
| PC 32:1 | [M+H]<sup>+</sup> | 732.554 | 732.5538 | 0.31 |
| PC 32:0/PC 16:0 16:0 | [M+H]<sup>+</sup> | 734.5697 | 734.5694 | 0.46 |
| PE 34:2/PE 16:0 18:2 | [M+Na]<sup>+</sup> | 738.5073 | 738.5045 | 3.85 |
| PE 36:5/PE 18:2 18:3 | [M+H]<sup>+</sup> | 738.5073 | 738.5068 | 0.73 |
| PC O-34:5 | [M+H]<sup>+</sup> | 738.5455 | 738.5432 | 3.07 |
| PC 32:1 | [M+Na]<sup>+</sup> | 754.5371 | 754.5357 | 1.84 |
| PC 34:4 | [M+H]<sup>+</sup> | 754.5371 | 754.5381 | -1.31 |
| PC 32:0/PC 16:0 16:0 | [M+Na]<sup>+</sup> | 756.5507 | 756.5514 | -0.83 |
| PC 34:3 | [M+H]<sup>+</sup> | 756.5507 | 756.5538 | -4.03 |
| PC 32:0/PC 16:0 16:0 | [M+Na]<sup>+</sup> | 756.5539 | 756.5514 | 3.36 |
| PC 34:3 | [M+H]<sup>+</sup> | 756.5539 | 756.5538 | 0.17 |
| PC 34:2/PC 16:0 18:2 | [M+H]<sup>+</sup> | 758.5696 | 758.5694 | 0.26 |
| PC 34:2/PC 16:0 18:2 | [M+H]<sup>+</sup> | 758.5731 | 758.5694 | 4.84 |
| PE 36:5/PE 18:2 18:3 | [M+Na]<sup>+</sup> | 760.4863 | 760.4887 | -3.20 |
| PE 36:5/PE 18:2 18:3 | [M+Na]<sup>+</sup> | 760.4908 | 760.4887 | 2.70 |
| PE 38:8 | [M+H]<sup>+</sup> | 760.4908 | 760.4912 | -0.58 |
| PC 34:1 | [M+H]<sup>+</sup> | 760.5817 | 760.5851 | -4.42 |
| PC 34:1 | [M+H]<sup>+</sup> | 760.5854 | 760.5851 | 0.41 |
| PG 34:4/PG 16:1 18:3 | [M+Na]<sup>+</sup> | 765.4674 | 765.4683 | -1.14 |
| PE 36:4/PE 18:2 18:2 | [M+K]<sup>+</sup> | 778.4772 | 778.4784 | -1.55 |
| PC 34:3 | [M+Na]<sup>+</sup> | 778.5359 | 778.5357 | 0.30 |
| PC 36:6/PC 18:3 18:3 | [M+H]<sup>+</sup> | 778.5359 | 778.5381 | -2.76 |
| PC 34:3 | [M+Na]<sup>+</sup> | 778.5394 | 778.5357 | 4.71 |
| PC 36:6/PC 18:3 18:3 | [M+H]<sup>+</sup> | 778.5394 | 778.5381 | 1.67 |
| PC 34:2/PC 16:0 18:2 | [M+Na]<sup>+</sup> | 780.5493 | 780.5514 | -2.59 |
| PC 34:2/PC 16:0 18:2 | [M+Na]<sup>+</sup> | 780.5525 | 780.5514 | 1.54 |
| PC 36:5 | [M+H]<sup>+</sup> | 780.5525 | 780.5538 | -1.56 |
| PC 36:5 | [M+H]<sup>+</sup> | 780.5557 | 780.5538 | 2.46 |
| PC 34:1 | [M+Na]<sup>+</sup> | 782.5691 | 782.567 | 2.66 |
| PC 36:4/PC 18:2 18:2 | [M+H]<sup>+</sup> | 782.5691 | 782.5694 | -0.40 |
| PC 36:4/PC 18:2 18:2 | [M+H]<sup>+</sup> | 782.5719 | 782.5694 | 3.22 |
| TG 44:3\|TG 8:0 18:0 18:3 | [M+K]+ | 783.5906 | 783.5905 | 0.14 |
| PE 38:7 | [M+Na]+ | 784.4872 | 784.4887 | -1.86 |
| PC 34:0 | [M+Na]+ | 784.5805 | 784.5827 | -2.77 |
| PC 34:0 | [M+Na]+ | 784.585 | 784.5827 | 3.06 |
| PC 36:3 | [M+H]+ | 784.585 | 784.5851 | -0.04 |
| PC 36:2 | [M+H]+ | 786.5973 | 786.6007 | -4.28 |
| PC 36:2 | [M+H]+ | 786.6008 | 786.6007 | 0.08 |
| PC 36:2 | [M+H]+ | 786.6046 | 786.6007 | 4.97 |
| PC 36:1 | [M+H]+ | 788.6135 | 788.6164 | -3.66 |
| PC 34:2\|PC 16:0 18:2 | [M+K]+ | 796.5235 | 796.5253 | -2.22 |
| PC 36:5 | [M+Na]+ | 802.5341 | 802.5357 | -2.04 |
| PC 36:5 | [M+Na]+ | 802.5379 | 802.5357 | 2.67 |
| PC 36:4\|PC 18:2 18:2 | [M+Na]+ | 804.5512 | 804.5514 | -0.20 |
| PC 36:4\|PC 18:2 18:2 | [M+Na]+ | 804.5548 | 804.5514 | 4.29 |
| PC 36:3 | [M+Na]+ | 806.5667 | 806.567 | -0.38 |
| PC 38:6 | [M+H]+ | 806.5667 | 806.5694 | -3.34 |
| PC 36:2 | [M+Na]+ | 808.5831 | 808.5827 | 0.61 |
| PC 36:1 | [M+Na]+ | 810.5978 | 810.5983 | -0.64 |
| TG 46:3\|TG 12:0 16:0 18:3 | [M+K]+ | 811.6224 | 811.6218 | 0.76 |
| TG O-48:2\|TG O-18:0_14:1_16:1 | [M+Na]+ | 811.7144 | 811.7155 | -1.40 |
| PC 35:1 | [M+K]+ | 812.5595 | 812.5566 | 3.56 |
| TG 50:3\|TG 16:0 16:0 18:3 | [M+Na]+ | 851.7094 | 851.7105 | -1.19 |
| TG 52:6\|TG 16:0 18:3 18:3 | [M+H]+ | 851.7094 | 851.7129 | -4.04 |
| TG 50:2\|TG 16:0 16:0 18:2 | [M+Na]+ | 853.722 | 853.7261 | -4.83 |
| TG 50:2\|TG 16:0 16:0 18:2 | [M+Na]+ | 853.7264 | 853.7261 | 0.34 |
| TG 52:5\|TG 16:0 18:2 18:3 | [M+H]+ | 853.7264 | 853.7285 | -2.45 |
| PI 34:2 | [M+Na]+ | 857.5158 | 857.5156 | 0.23 |
| PI 36:5 | [M+H]+ | 857.5158 | 857.518 | -2.53 |
| TG 52:9:1O\|TG 16:3 18:3 18:3:1O | [M+H]+ | 861.6587 | 861.6608 | -2.44 |
| TG 52:7:1O\|TG 18:2 18:3 16:2:1O | [M+H]+ | 865.6909 | 865.6921 | -1.40 |
| TG 52:6\|TG 16:0 18:3 18:3 | [M+Na]+ | 873.6955 | 873.6948 | 0.72 |
| TG 54:9\|TG 18:3 18:3 18:3 | [M+H]+ | 873.6955 | 873.6972 | -1.99 |
| TG 54:9\|TG 18:3 18:3 18:3 | [M+H]+ | 873.6999 | 873.6972 | 3.08 |
| TG 52:5\|TG 16:0 18:2 18:3 | [M+Na]+ | 875.7095 | 875.7105 | -1.07 |
| TG 54:8\|TG 18:2 18:3 18:3 | [M+H]+ | 875.7095 | 875.7129 | -3.84 |
| TG 52:5\|TG 16:0 18:2 18:3 | [M+Na]+ | 875.714 | 875.7105 | 4.00 |
| TG 54:8\|TG 18:2 18:3 18:3 | [M+H]+ | 875.714 | 875.7129 | 1.22 |
| TG 52:4\|TG 16:0 18:2 18:2 | [M+Na]+ | 877.7247 | 877.7261 | -1.60 |
| TG 54:7\|TG 18:2 18:2 18:3 | [M+H]+ | 877.7247 | 877.7285 | -4.31 |
| TG 52:4\|TG 16:0 18:2 18:2 | [M+Na]+ | 877.7289 | 877.7261 | 3.11 |
| TG 54:7\|TG 18:2 18:2 18:3 | [M+H]+ | 877.7289 | 877.7285 | 0.41 |
| TG 52:3\|TG 16:0 18:1 18:2 | [M+Na]+ | 879.7402 | 879.7418 | -1.80 |
| TG 54:6\|TG 18:2 18:2 18:2 | [M+H]+ | 879.7402 | 879.7442 | -4.55 |
| TG 52:2\|TG 16:0 18:0 18:2 | [M+Na]+ | 881.7562 | 881.7575 | -1.43 |
| TG 54:5\|TG 18:0 18:2 18:3 | [M+H]+ | 881.7562 | 881.7598 | -4.12 |
| TG 52:2\|TG 16:0 18:0 18:2 | [M+Na]+ | 881.7597 | 881.7575 | 2.56 |
| TG 54:5\|TG 18:0 18:2 18:3 | [M+H]+ | 881.7597 | 881.7598 | -0.12 |
| PI 36:2 | [M+Na]+ | 885.5487 | 885.5469 | 2.03 |
| TG 52:6\|TG 16:0 18:3 18:3 | [M+K]+ | 889.6696 | 889.6688 | 0.98 |
| TG 52:5\|TG 16:0 18:2 18:3 | [M+K]+ | 891.6864 | 891.6844 | 2.27 |
| TG 54:9\|TG 18:3 18:3 18:3 | [M+Na]+ | 895.679 | 895.6792 | -0.15 |
| TG 54:8\|TG 18:2 18:3 18:3 | [M+Na]+ | 897.6958 | 897.6948 | 1.11 |
| TG 54:7\|TG 18:2 18:2 18:3 | [M+Na]+ | 899.709 | 899.7105 | -1.67 |
| TG 54:6\|TG 18:2 18:2 18:2 | [M+Na]+ | 901.7254 | 901.7261 | -0.79 |
| TG 54:4\|TG 18:0 18:2 18:2 | [M+Na]+ | 905.7585 | 905.7575 | 1.16 |
| TG 54:6\|TG 18:2 18:2 18:2 | [M+K]+ | 917.6982 | 917.7001 | -1.99 |
| TG 54:5:1O\|TG 18:0 18:2 18:3:1O | [M+K]+ | 935.7141 | 935.7106 | 3.79 |
| TG 54:5:2O\|TG 18:0 16:3 20:2:2O | [M+Na]+ | 935.7272 | 935.7315 | -4.61 |
| TG 55:4\|TG 18:1 19:1 18:2 | [M+K]+ | 935.7425 | 935.747 | -4.74 |
| TG 56:3\|TG 16:0 22:0 18:3 | [M+Na]+ | 935.8028 | 935.8043 | -1.65 |
| TG 58:6\|TG 22:0 18:3 18:3 | [M+H]+ | 935.8028 | 935.8068 | -4.31 |
| TG 60:2\|TG 16:0 26:0 18:2 | [M+H]+ | 971.9038 | 971.9007 | 3.18 |
| TG 60:1\|TG 16:0 26:0 18:1 | [M+H]+ | 973.9156 | 973.9163 | -0.78 |

Note: the m/z Value depicts the measured mass via MALDI MS, while m/z Match is the LCMS/MS-matched value. The mass error for the measured masses compared to the exact masses are represented by PPM error (parts per million).

